# Encoding of continuous perceptual choices in human early visual cortex

**DOI:** 10.1101/2023.02.10.527876

**Authors:** Riccardo Barbieri, Felix M. Töpfer, Joram Soch, Carsten Bogler, Henning Sprekeler, John-Dylan Haynes

## Abstract

Research on the neural mechanisms of perceptual decision-making has typically focused on simple categorical choices, say between two alternative motion directions. Studies on such discrete alternatives have often suggested that choices are encoded either in a motor-based or in an abstract, categorical format in regions beyond sensory cortex. However, many sensory features are graded rather than discrete, raising the question how choices are encoded when they span the full sensory continuum. Here we assessed this using motion stimuli that could vary anywhere between 0° and 360°. We employed a combination of neuroimaging and encoding models based on Gaussian Process Regression to assess how either stimuli or choices were encoded in brain responses. We found that single-voxel tuning patterns could be used to reconstruct the trial-by-trial physical direction of motion as well as the participants’ continuous choices. Importantly, these continuous choice signals were primarily observed in early visual areas. The tuning properties in this region generalized between choice encoding and stimulus encoding, even for reports that reflected pure guessing. We found only little information related to the decision outcome in regions beyond visual cortex, such as parietal cortex, possibly because our task did not involve differential motor preparation. This could suggest that decisions for continuous stimuli take can place already in sensory brain regions, potentially using similar mechanisms to the sensory recruitment in visual working memory.

## INTRODUCTION

The brain mechanisms of perceptual decisions involve several sequential steps (Gold & Shadlen, 2007): First, information about an external stimulus is encoded in sensory brain regions (e.g. whether an object is moving leftward, or rightward). Then, the sensory evidence supporting the different potential states of the world is gathered in a decision variable, which typically also integrates information over time (Gold & Shadlen, 2007; but see Uchida et al., 2006). Finally, once enough evidence is collected in favor of a certain hypothesis, a decision is made and the observer takes an action to indicate their choice.

Several stages of this processes have been characterized in detail, both in humans and in monkeys (Gold & Shadlen, 2007; Heekeren et al., 2008; Mulder et al., 2014; Forstmann et al., 2016; Hanks & Summerfield, 2017). In monkeys, sensory neurons encode stimulus-related information, such as the physical direction of movement of dots in the visual field (Shadlen et al., 1996). Regions of parietal and frontal cortex encode the gradual accumulation of choice-related, categorical evidence in monkeys (Gold & Shadlen, 2007) and in humans (Siegel et al., 2011; Mulder et al., 2014; Wilming et al., 2020). Neuroimaging signals also reflect levels of evidence (Heekeren et al., 2004; Forstmann et al., 2016) and strategic adjustments in decisions (Mulder et al., 2014). As for the outcome of the decision, many human neuroimaging studies have identified choice-related brain signals in several areas including parietal cortex (Tosoni et al., 2008, 2014; Liu & Pleskac, 2011; Hebart et al., 2012, 2016; Levine & Schwarzbach, 2017), insular cortex (Liu & Pleskac, 2011; Ho et al., 2009) and prefrontal cortex (Heekeren et al., 2004, 2006; Filimon et al., 2013; Hebart et al., 2016). Interestingly, studies have shown that activity already in sensory areas is influenced by the behavioral choice, especially in ambiguous stimulus conditions (Ress & Heeger, 2003; Serences & Boynton, 2007; Sousa et al., 2021).

Due to this variety of decision signals, the representational space in which decision outcomes are encoded has remained somewhat unclear, with some suggesting a motor-based “intentional” frame of reference (Shadlen et al., 2008; Tosoni et al., 2014) whereas others have shown that decision signals can be dissociated from motor plans in both monkeys and humans (Bennur & Gold, 2011; Hebart et al., 2012; Filimon et al., 2013; Park et al., 2014; Brincat et al., 2018, but see discussion below).

Importantly, there is an issue that has only received little attention: Studies of perceptual decision making have typically employed few discrete alternative stimulus features, whereas perception of most sensory features is inherently continuous (Levinson & Sekuler, 1976; Nichols & Newsome, 2002; Prinzmetal et al., 1998; van Bergen et al., 2015). For example, most studies that use perceptual judgements of coherent motion focus on few alternative motion directions in each trial and require categorical responses (Gold & Shadlen, 2007; Huk & Meister, 2012; Hanks & Summerfield, 2017). In these tasks a small set of possible motion directions (e.g. motion left or motion right) has to be mapped onto a small set of predetermined motor responses (e.g. pressing the left or the right button or making a saccade to a left or right target). In such cases, participants might encode their choices in a motor frame of reference or in a lower-dimensional categorical form (“left” vs “right”). In contrast, choices could also be encoded in some kind of continuous perceptual space (Beck et al. 2008; van Bergen et al., 2015; Smith, 2016; Ratcliff, 2018). Note that even if a paradigm allows to dissociate choices from motor plans by the use of trial-wise varying stimulus-response mappings (Bennur & Gold, 2011; Hebart et al. 2012), the encoding of choices still occurs in the form of such stimulus-response-mappings and thus uses discrete, lower-dimensional representations. Alternatively, however, choices could be encoded on a full 360° continuum.

## RESULTS

Here we assessed the encoding of continuous perceptual choices using a combination of fMRI and voxel-wise fMRI encoding models (Nevado et al., 2004; Thirion et al., 2006; Kay et al., 2008; Dumoulin et al., 2008; Brouwer & Heeger, 2009; Naselaris et al., 2011; Haynes, 2015; see Materials and Methods for full details). As stimuli we used random dot kinematograms (RDKs) as in many studies of perceptual decision making (Newsome & Parè, 1988). These stimuli consist of an array of dots moving in various directions like a detuned TV-set. By modifying the proportion of dots that coherently move in a single target direction (signal) among others moving in random directions (noise), it is possible to assess perceptual decisions under varying levels of sensory information.

For our feature-continuous motion stimuli the directions were drawn from a uniform distribution between 0° and 360°. Participants reported their judgements by pressing a button when a rotating sensory comparison stimulus matched their choice (Figure 1). In previous work we found that reports like this that used a sensory reference stimulus, instead of e.g. the movement of a track ball, had the highest accuracy for continuous judgements (Töpfer et al., 2022). It also allows to decouple choice-related signals from specific motor preparation. We measured trial-by-trial brain activity under three different coherence levels: 0%, intermediate and 100% coherence. Note that at 0% coherence there is no physical evidence regarding the stimulus direction and participants are purely guessing. This condition is of particular importance because it allows to study choices independent of physical stimulus information, and it will be the primary focus of our preregistered analyses. In contrast, at 100% coherence stimulus direction and choices are highly correlated (see below).

**Figure 1:**
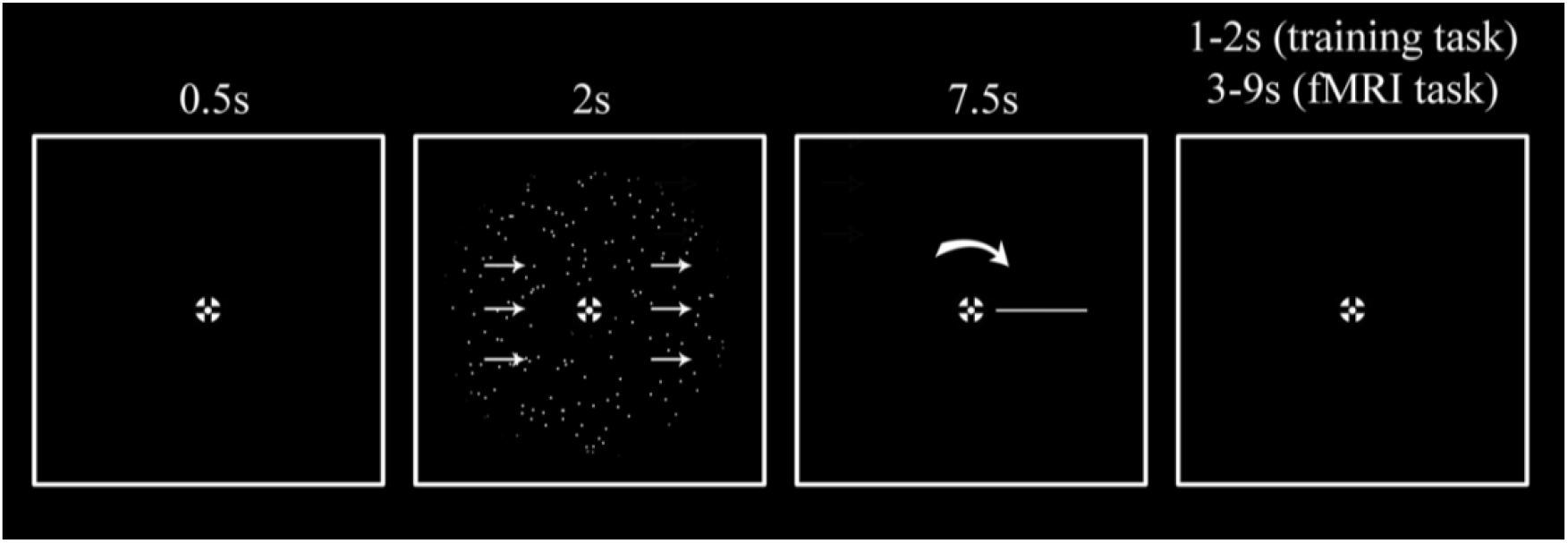
A single trial stimulus sequence. Participants were instructed to fixate the center of a bullseye for the entire duration of the experiment. 0.5 s after trial onset, they were presented with 2 s of random dot motion stimulus (RDK) displayed with a different direction of motion and coherence level for every trial. The direction of motion was continuously distributed and spanned between 0° and 360°. For the training task participants were presented with five different coherence levels (0%, 12.5%, 25%, 50% and 100%). For the fMRI task, instead, only three coherence levels were presented (0%, an intermediate coherence level individually estimated during the training phase, and 100%). Based on previous experiments (Töpfer et al., 2022) we used a report method that involved a rotating perceptual comparison stimulus and thus was given in a perceptual (as opposed to a motor) frame of reference. This has the advantage of increasing accuracy and minimizing bias (Töpfer et al., 2022). Specifically, participants were instructed to report the net motion direction by pressing the response button when a self-moving rotating bar on the screen matched the direction of motion they perceived. After the report was given the bar continued its cycle so that the presentation duration was always 7.5 s. Inter-trial-intervals (ITIs) for the training task were 1 or 2s, for the fMRI task they were between 3s, 5s, 7s and 9s, exponentially distributed so that shorter ITIs happened more frequently than longer ones.

Participants were able to correctly perform the continuous task with increasing accuracy the higher the coherence of the stimuli (Figure 2). When sensory information was absent (0% coherence) there was no relationship between direction and report, thus as expected participants responded largely randomly (Figure 2 A,B left panels). With increasing sensory evidence the distributions of responses became narrower and centered around the veridical direction (Figure 2 A,B mid and right panels). This was also reflected in increasing levels of accuracy with increasing coherence (Figure 2C). As expected from previous work (Töpfer et al., 2022) our chosen report method (see Methods) minimized the reporting biases across the motion directions.

**Figure 2:**
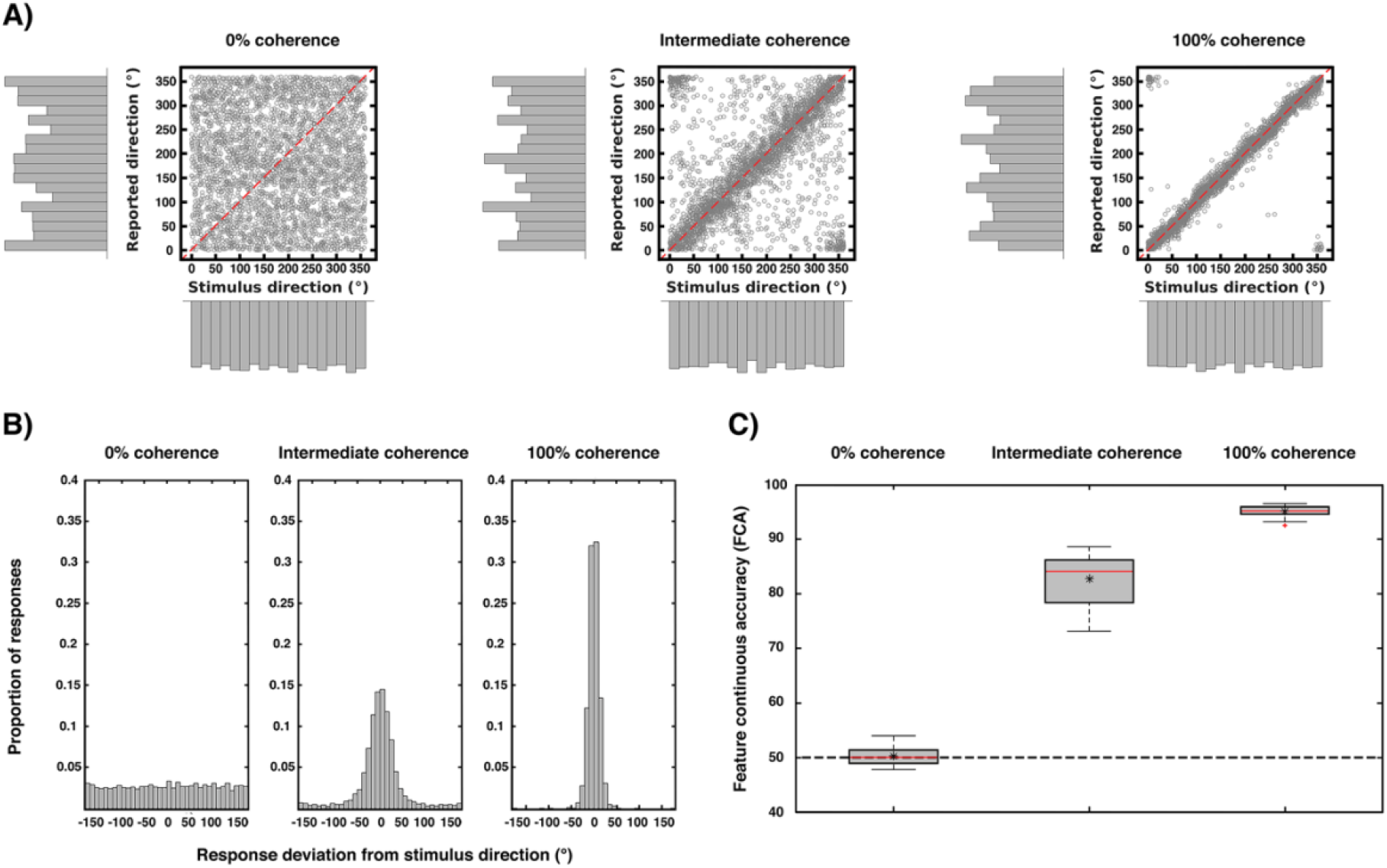
Behavioral results. The three plots show the behavioral performance pooled from all subjects (N=23) for each coherence level. **(A)** Scatterplots and marginal distributions of the stimulus and reported directions for each level of coherence (each point corresponds to one trial). **(B)** Distribution of participants’ response deviations (where 0° corresponds to perfect performance). Note the flat distribution when direction information is absent (0% coherence) and the narrowing of the response distributions with increasing coherence. **(C)** Performance accuracy (measured as deviations as in **B** but replotted as FCA, which is scaled such that 100 is perfect performance and 50 is chance performance; see Methods). In each box, the central red line indicates the median accuracy, the asterisk indicates the mean accuracy. The bottom and top edges of the box indicate the 25th and 75th percentiles, respectively. The whiskers extend to the most extreme data points not considered outliers, and the outliers are plotted individually using a red cross. The dashed black line indicates chance performance. There was a main effect of coherence on accuracy (repeated measures ANOVA; F(1.26, 27.818) = 1357.545; p < 0.001; Greenhouse-Geisser corrected for non-sphericity).

The brain signals associated with these behavioral judgements were then entered into two different preregistered encoding model analyses: One that assessed encoding of the *physical* stimulus direction, and the other that modelled the encoding of the participants’ trial-by-trial *reports*. Our model extends the framework of so-called choice probabilities that were primarily developed for binary choices (Britten et al., 1996; Chicharro et al., 2021) to continuous encoding. In comparison to other studies, our encoding models were based on a cyclic version of Gaussian Process Regression (GPR; Rasmussen & Williams, 2005; Caywood et al., 2017; Dimitrova et al., 2020). This has the advantage of providing not only an estimate of the mean response in each voxel for a given direction, but also of the distribution across trials, separately for each direction, allowing to obtain tuning response profiles (see Methods for details). Also, this approach does not require any a priori assumptions regarding the smoothness of the tuning functions (compared to e.g. Brouwer & Heeger, 2009). Figure 3 shows examples of these voxel tuning functions for sixteen visually responsive voxels of one participant. There is considerable variability with some voxels showing various forms of smoothly varying tuning functions, while others are non-informative and flat. Note that these tuning functions cannot be directly interpreted in terms of single-neuron tuning, but reflect a complex integration of a population of tuned neurons within a voxel (Kriegeskorte et al., 2010; Ramirez et al., 2014; Sprague et al., 2018; Gardner & Liu, 2019, Sprague et al., 2019).

**Figure 3:**
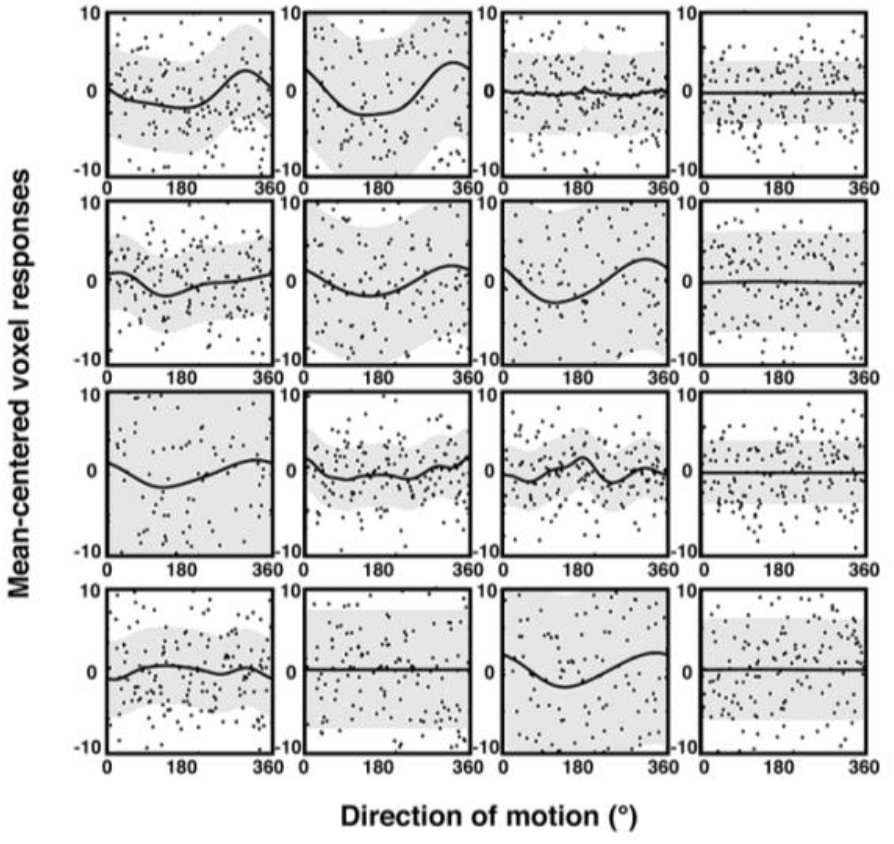
Trial-wise activity and estimated sensory voxel tuning response profiles based on GPR (see Methods) for encoding of physical stimulus direction. The plot shows the trial-wise fMRI response (black dots) as a function of motion direction, along with the estimated GPR-based tuning functions of 16 voxels (mean = black lines; standard deviation = gray bands; example participant; 100% coherence condition). The voxels were randomly chosen from the subject’s most informative searchlight of the ROI described in the exploratory analyses (see *Additional exploratory analyses* in the Materials and Methods section). Trial-wise activity (y-axis) is here plotted as parameter estimates of a trial-wise 1^st^ level GLM versus corresponding stimulus motion directions (x-axis). The estimated response profiles (thick lines, grey regions) were obtained by averaging the voxel-wise GPR predictions across 10 cross-validation folds. The data were mean-centered and are shown at the same scale to facilitate comparison.

In the next step, we used ensembles of voxel-wise tuning functions from different pre-defined ROIs to reconstruct (a) the veridical motion direction and (b) the reported direction in each given trial (see Figures 4 and 5 for details on methods). These five bilateral ROIs were early visual cortex (EVC), MT+, superior parietal cortex (SPC), intraparietal sulcus (IPS), and inferior parietal cortex (IPC). These areas have been reported in previous work to be involved in encoding of motion and decision signals (Kamitani & Tong, 2006; Serences & Boynton, 2007; Hebart et al, 2012; Bode et al., 2013; Levine & Schwarzbach, 2017).

**Figure 4:**
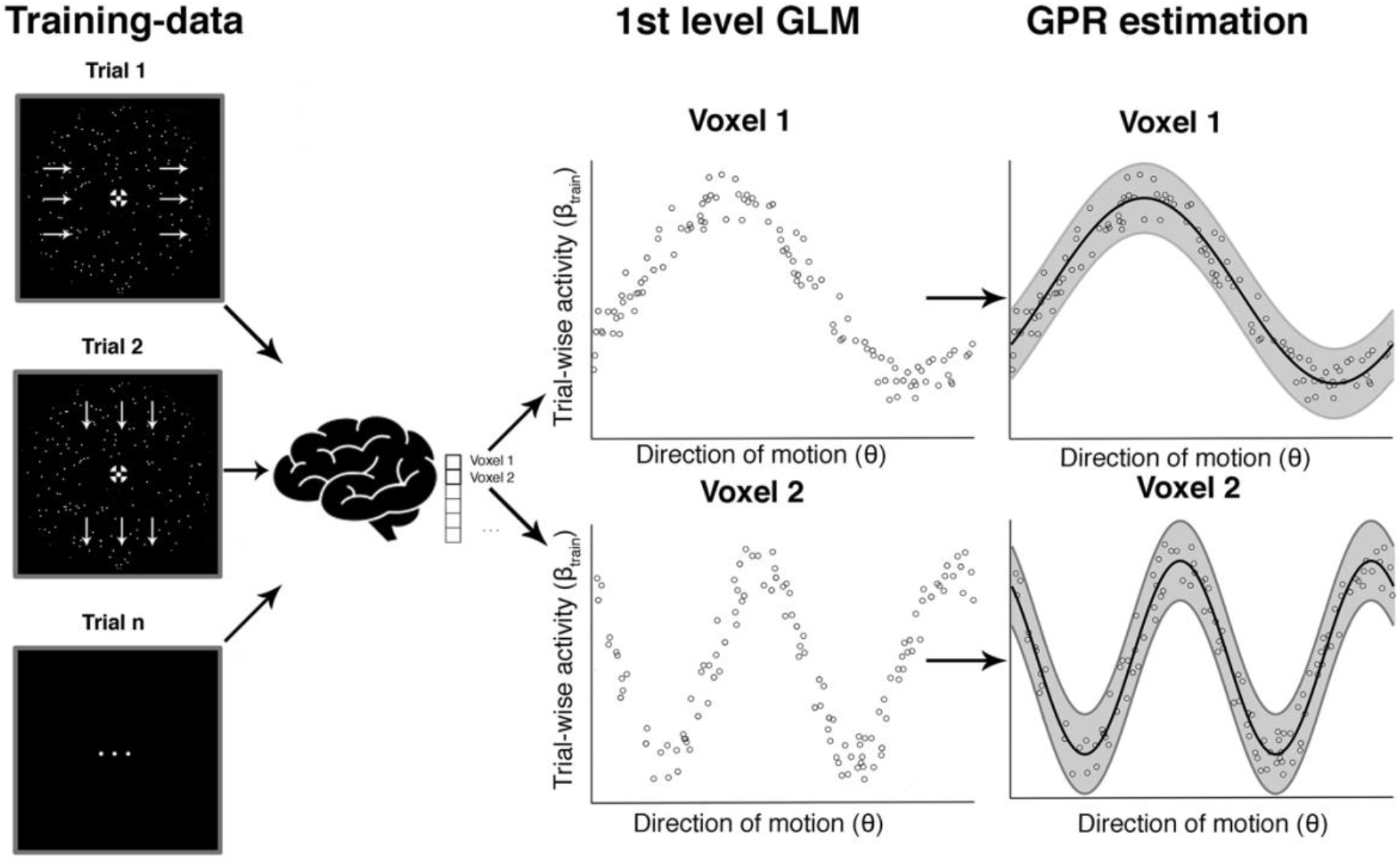
*Voxel-wise GPR estimation to obtain tuning response profiles*. This schematic example is based on simulated data and illustrates the steps involved in the voxel-wise GPR estimation. For a given direction of motion in a given trial we obtain a (trial-wise) parameter estimate for the response amplitude in each voxel. The middle shows a set of trial-wise measurements for two example voxels. The x-axis corresponds to the direction of motion in a trial and the y-axis to the trial-wise activity in that voxel. Each vertical section is equivalent to a likelihood function of a brain response amplitude in a voxel given a certain stimulus direction. We use Gaussian Process Regression (GPR) to estimate the family of these likelihood functions across different movement directions (right column, “GPR estimation”). The analysis can be performed either with the veridical stimulus motion direction *θ*_*s*_ or the reported direction *θ*_*r*_.

**Figure 5:**
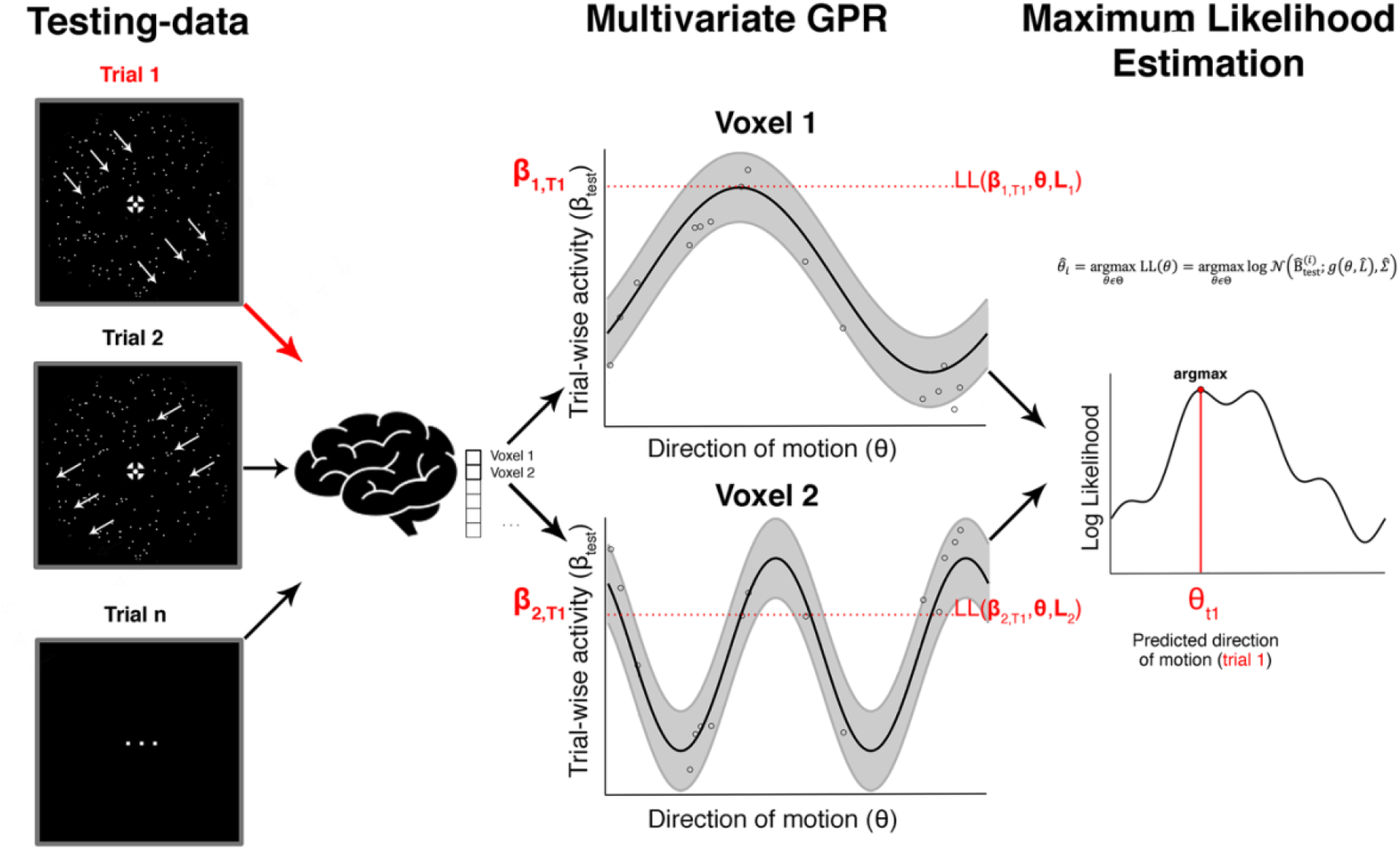
*Searchlight-based GPR reconstruction from a multivoxel ensemble of tuning response profiles*. This shows the steps involved in the searchlight-based reconstruction of one trial focusing on just two voxels. The red arrow on the left shows an unknown true direction on a single trial T_1_. The aim is to estimate this true direction with maximum precision from an ensemble of trial-wise brain responses using the tuning response profiles. The red parameters on the y-axis in the middle column (β_1, T1_ and β_2, T1_) show the measured brain responses in that trial in the two voxels. The horizontal red section illustrates the likelihood values for each direction. If one were only using a single voxel one would estimate the direction as that where the likelihood is maximal (assuming a flat prior as in our study). In our case we combine the likelihood estimates across an ensemble of voxels (in a searchlight) and identify the maximal value, which corresponds to our decoded direction of motion in that trial.

Figure 6 shows the stimulus-related and report-related accuracies for each of these ROIs and each coherence level. For each ROI, we tested whether coherence or label (stimulus versus report) influenced accuracy. Based on previous research on motion direction decoding with fMRI we expected that fMRI-signals would carry highest information in early visual cortex (Kamitani & Tong, 2006; Serences & Boynton, 2007; Hebart et al, 2012), whereas it was unclear whether this region is expected to encode choice (Ress & Heeger 2003; Serences & Boynton, 2007; Brouwer & van Ee 2007; Krishna et al., 2021). As expected, we found a main effect of coherence on reconstruction performance for the visual ROI (coherence on EVC: F(2, 44) = 11.909, p < 0.001) but there was no difference between accuracies for stimuli and reports and no interaction (F(1,22) = 2.42, p = 0.134; label*coherence F(2,44) = 0.69, p = 0.507). A post-hoc t-test performed on EVC revealed no difference between the stimulus and the report reconstruction at 0% coherence (t = -1.793; p = 1, Bonferroni corrected for a family of 15 multiple comparisons).

**Figure 6:**
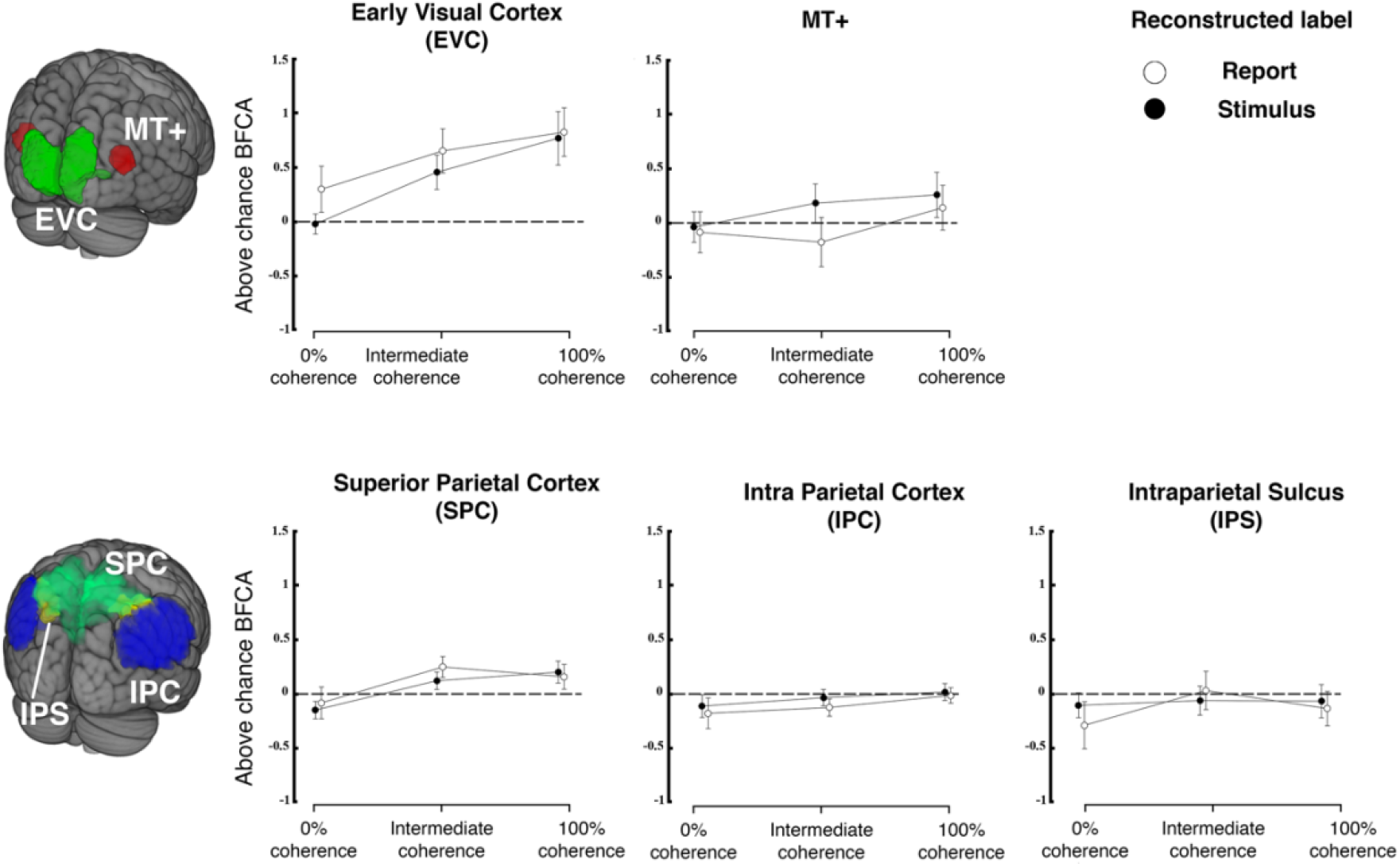
Reconstruction from regions of interest in visual and parietal cortex. The top row shows reconstruction accuracy (expressed as balanced feature continuous accuracy, BFCA, which is corrected for biases due to unequal distribution of responses across the sensory continuum, see Methods) for regions of interest in early visual cortex (comprising V1, V2 and V3) and MT+ separately for each coherence level. The bottom row shows the same for three regions of parietal cortex. Filled versus open symbols indicate reconstruction of stimulus and report respectively. Error bars are standard errors. Dashed lines indicate chance level (N = 23).

As in some (but not all) previous studies we also expected the motion sensitive complex MT+ to be informative of veridical motion direction (Kamitani & Tong, 2006) and of perceptual judgements (Serences & Boynton, 2007). We also expected a main effect of coherence on stimulus and report reconstruction (Britten et al., 1996). In contrast to this prediction, the results of a repeated measures ANOVA revealed no effect of coherence in MT+ (F(2, 44) = 0.664, p = 0.52), no effect of which label was being reconstructed (label on MT+: F(1,22) = 2.72, p = 0.113) and no interaction (label * coherence on MT+: F(2,44) = 0.628, p = 0.538). Additional post-hoc t-tests revealed that the mean stimulus reconstruction performance at 100% coherence was not different from that of 0% coherence, in which the stimulus has no net direction of motion (t = 1.069; p = 1, Bonferroni corrected for a family of 15 multiple comparisons). Please note that these differences to previous work might reflect the fact that we employed continuous stimuli and also that we used a very different response format that did not involve differential or even dispositional motor preparation.

We also found a main effect of coherence on reconstruction performance for one region of parietal cortex (SPC; coherence SPC: F(2,44) = 4.219, p = 0.021), but no effect of which label was reconstructed (label SPC: F(1,22) = 0.712, p = 0.408) and no interactions were found (label*coherence SPC: F(2,44) = 0.502, p = 0.609). We did not find any effect on reconstruction performance for the other parietal ROIs (IPS and IPC).

We then conducted an exploratory analysis aimed to identify informative regions beyond our pre-defined regional hypotheses. For this, we used whole-brain searchlight reconstruction maps (see Methods). In line with our ROI-based findings this only revealed a cluster in early visual cortex (left occipital pole; Appendix 1 – Figure 2). Interestingly, we found no region where coherence affected reconstruction of choices (see Appendix 1 - *Statistical analyses: searchlight-based reconstructions and effect of coherence*).

Taken together, the early visual cortex is the only area where we were able to reconstruct information about continuous motion stimuli and their corresponding choices. At 0% coherence we found no evidence for either stimulus encoding (as expected) or response encoding in our early visual ROI. In constrast, other studies have revealed choice signals already in early visual areas (Ress & Heeger, 2003; Serences & Boynton 2007; Sousa et al. 2021). We thus assessed whether our a priori defined early visual ROI might be defined too widely to reveal potential differences between stimulus and report reconstruction. For this, we conducted a single test with a more focused analysis of the voxels that encoded both stimulus and report across all of the coherence levels (0%, Intermediate and 100%). Such voxels were located in the occipital cortex bilaterally. We found a label*coherence interaction effect on the reconstruction performance in these areas (F(1.327, 29.191) = 4.426, p = 0.034; Greenhouse-Geisser correction for non-sphericity; see Figure 7). Moreover, post-hoc t-tests revealed a significant difference between stimulus and report reconstruction at 0% coherence for this more focused voxel set (Table 1 - t = 3.560; p = 0.011, Bonferroni corrected for a family of 15 multiple comparisons). Please note that the voxel selection for this analysis was collapsed across both stimulus and report reconstruction and was thus not biased a priori to yield such a difference.

**Figure 7:**
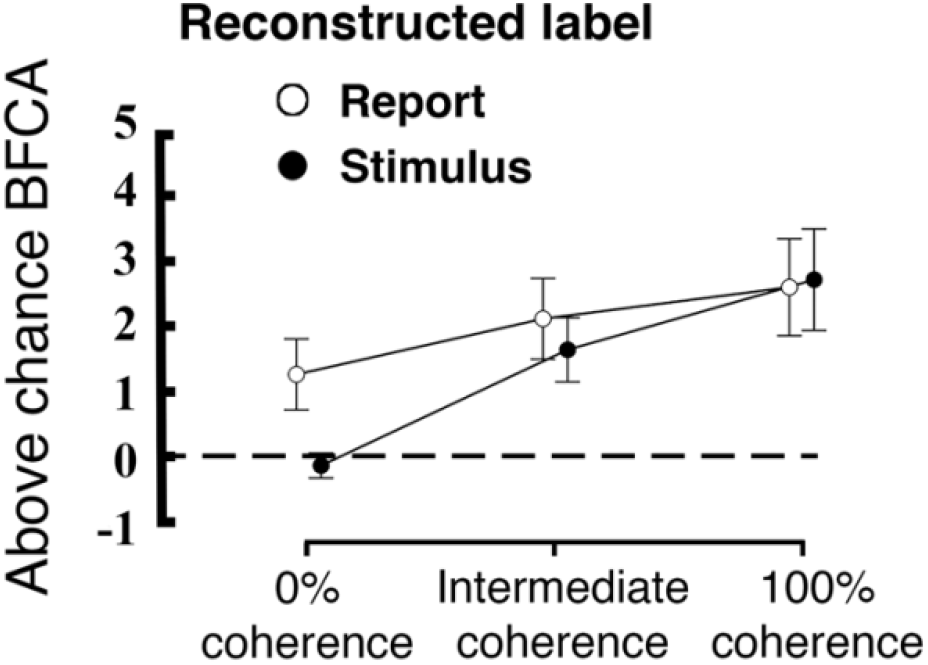
Interaction between coherence level and reconstructed label within a subsection of the early visual region (the conjunction of voxels that code for both stimulus and report). The plot displays the averaged above-chance accuracy (expressed as BFCA, see Methods) across voxels for each coherence level and each reconstructed label, error bars are standard errors, the black dashed line represent chance level (N = 23). Please note that this voxel selection does not affect the difference between encoding of stimulus versus reports.

**Table 1:** Post-hoc t-tests performed on the stimulus and report reconstruction performance. Please, note that in the 0% coherence condition, the stimulus labels are unrelated with the motion direction (motion signal is absent, and labels are randomly assigned), hence the chance-level reconstruction performance is to be expected.

|  |  | Mean Difference | SE | t | P<sub>bonf</sub> |
| --- | --- | --- | --- | --- | --- |
| Coherence 0%, Report | Coherence Intermediate, Report | -0.854 | 0.508 | -1.682 | 1.000 |
|  | Coherence 100%, Report | -1.338 | 0.508 | -2.635 | 0.155 |
|  | Coherence 0%, Stimulus | 1.397 | 0.392 | 3.560 | 0.011 * |
|  | Coherence Intermediate, Stimulus | -0.379 | 0.530 | -0.715 | 1.000 |
|  | Coherence 100%, Stimulus | -1.453 | 0.530 | -2.744 | 0.114 |
| Coherence Intermediate, Report | Coherence 100%, Report | -0.484 | 0.508 | -0.953 | 1.000 |
|  | Coherence 0%, Stimulus | 2.251 | 0.530 | 4.250 | < .001 *** |
|  | Coherence Intermediate, Stimulus | 0.475 | 0.392 | 1.211 | 1.000 |
|  | Coherence 100%, Stimulus | -0.599 | 0.530 | -1.131 | 1.000 |
| Coherence 100%, Report | Coherence 0%, Stimulus | 2.735 | 0.530 | 5.164 | < .001 *** |
|  | Coherence Intermediate, Stimulus | 0.959 | 0.530 | 1.810 | 1.000 |
|  | Coherence 100%, Stimulus | -0.115 | 0.392 | -0.294 | 1.000 |
| Coherence 0%, Stimulus | Coherence Intermediate, Stimulus | -1.776 | 0.508 | -3.498 | 0.012 * |
|  | Coherence 100%, Stimulus | -2.850 | 0.508 | -5.613 | < .001 *** |
| Coherence Intermediate, Stimulus | Coherence 100%, Stimulus | -1.074 | 0.508 | -2.116 | 0.568 |

Note. P-value adjusted for comparing a family of 15
\* p < .05, \*\* p < .01, \*\*\* p < .001

### Model generalization

In order to further test these choice-related signals, we performed a specifically targeted cross-prediction analysis. We took the report-related reconstruction model estimated at the 0% coherence condition and used it to cross-predict the stimuli in the 100% coherence condition. This model generalization constitutes an independent test of the choice signals. Importantly, by training the reconstruction model on the 0% condition we avoid that our model is influenced by residual sensory information. This tests whether the choices at 0% coherence, i.e. when participants are guessing, use a similar representational format as the encoding of stimuli. Indeed, the 0% coherence report model generalized to the 100% stimulus condition (and vice-versa) (right tailed one sample t-test; t(22) = 2.969; p = 0.004). For exploratory purposes we also repeated this procedure with stimulus motion directions in the intermediate and 0% coherence condition. The generalization tests between the different reporting conditions constitute a test of model consistency across evidence levels and were all above chance. As predicted the cross-prediction was significant between 0% coherence report and the two above-chance stimulus encoding conditions, but not the 0% coherence condition where there is no sensory information. The summary of this cross-prediction analysis is displayed in Figure 8. Thus, we conclude that choice-related signals are present at guessing levels in early visual cortex and that these signals are encoded in a similar form as the physical stimulus features.

**Figure 8:**
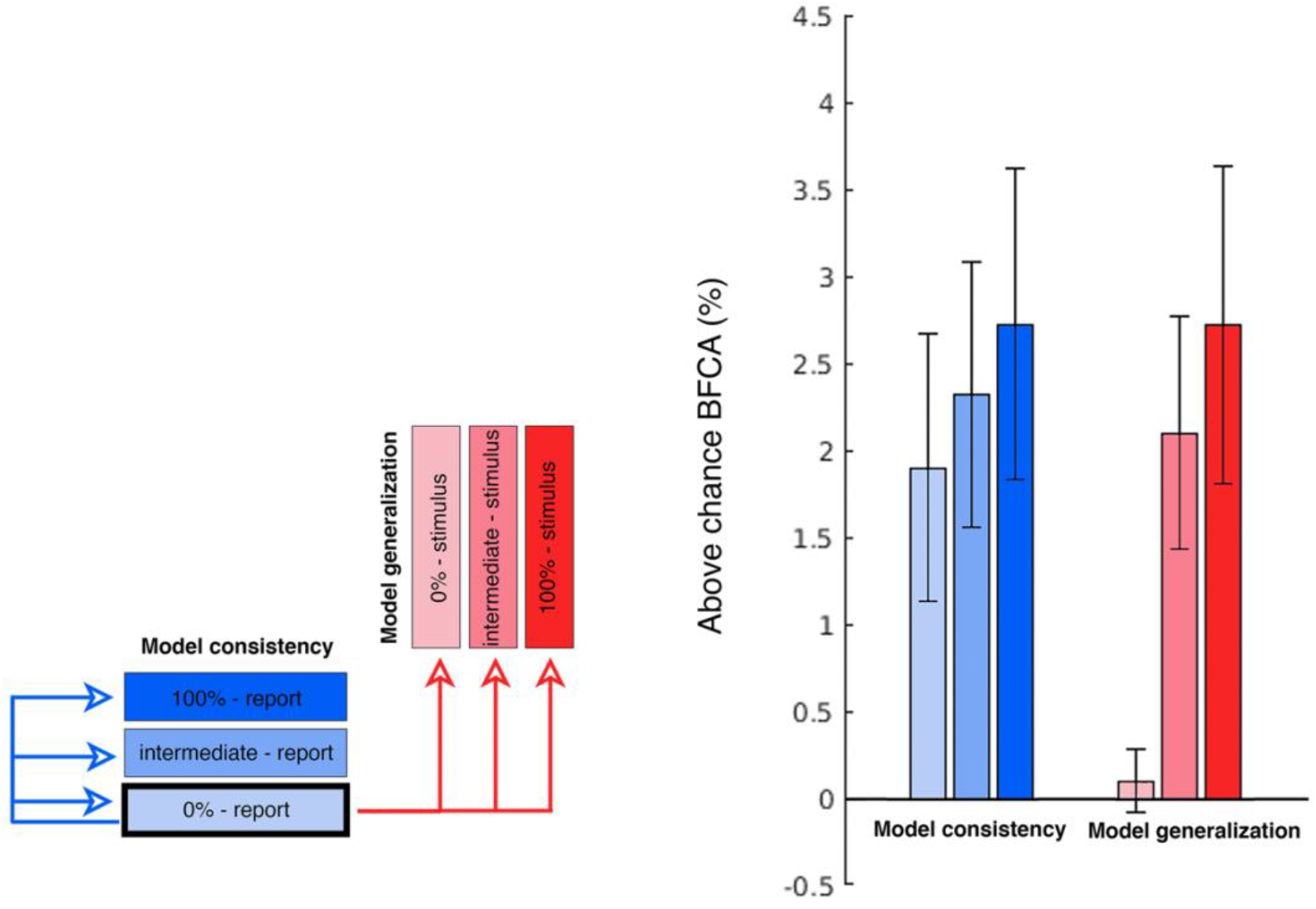
Cross-prediction performances. The figure summarizes the cross-prediction analyses performed to test the model generalization and model consistency. Each bar displays the above-chance generalization accuracy (expressed as BFCA, see Methods) of the 0% report report-related GPR model to every other condition. The bars on the right hand of the plot (in red) indicate how well the 0% report model allows to reconstruct the stimulus identity in each coherence condition (model generalization). We were primarily interested in observing the generalization performance of the 0% coherence report model to the 100% stimulus (first bar on the right). Also note that since the 0% coherence stimulus model is based on random directions (unrelated with the report or the stimulus), it represents here a form of control condition in which chance-level generalization performance is to be expected (the first red bar in the right group). The group of bars on the left hand of the plot (in blue) illustrates how well the 0% report model allow to reconstruct behavioral reports in each coherence condition (model consistency)(N=23; errorbars: SEM).

In order to control for involuntary eye movements that might have confounded our reconstruction analyses, we had first decided to exclude trials where subjects exceed a threshold of 2 dva for more than 200 ms during the stimulation period. However, recent evidence indicates that involuntary eye movements below our adopted threshold might still affect brain activity, thus constituting a potential confound (Merriam et al., 2013; Thielen et al., 2019). For this reason, we complemented our preregistered results with an additional analysis that exploited the same GPR-based estimation and reconstruction techniques adopted for the analysis of brain signals, to the eye movements recorded with eye-tracker (see Appendix 1 - *Control analysis: stimulus and report reconstruction from eye-tracking data*). Our results revealed that the pattern of eye movement was unrelated to the stimulus or the report in the 100% and the intermediate coherence condition, but it was predictive of participants’ reports in the 0% coherence condition (Appendix 1 – Figure 3). Thus, it could have been possible that the choice signals in the guessing condition were partly affected by eye movements. However, this would not explain why our report model generalized from the guessing to the other conditions, where eye movements did not play a role.

## DISCUSSION

Previous studies on neural mechanisms of perceptual decision making have often focused on simple decisions involving discrete alternatives. These have found discrete decisions to be encoded either in intention-related formats (Shadlen et al., 2008), or in high-level but effector-independent formats (Heekeren et al., 2008; Bennur & Gold, 2011; Hebart et al., 2012; Park et al., 2014; Brincat et al., 2018). However, it has remained unclear how the brain encodes perceptual choices regarding an entire continuum of features, which is what our study aimed to address (see also Nichols & Newsome, 2002; van Bergen et al., 2015; Ratcliff, 2018 for other examples of continuous stimuli). Our study also disentangled stimulus and choice-related activity from motor responses by using a visual comparison stimulus (a rotating bar). Our modelling of single-voxel responses as continuously varying distributions using Gaussian Process Regression (GPR; Rasmussen & Williams, 2005) allowed us to estimate tuning functions in considerable detail and without making a priori assumptions regarding their smoothness.

Our design revealed that the activity of visual voxels was modulated by both the stimulus directions across trials (Figure 3) and by participants’ perceptual choices regarding the motion direction. We were able to reconstruct the stimulus motion direction from clusters of voxels in early visual cortex (Figure 6), and to identify a region of occipital cortex in which the change in coherence level had a different effect on stimulus and report reconstruction (Figure 7). These findings extend previous research on decoding of visual motion direction and perceptual choices from brain signals (Kamitani & Tong, 2006, Serences & Boynton, 2007; Hebart et al, 2012) to the feature-continuous case. Moreover, they indicate that perceptual decisions can be represented by neural populations encoding the stimulus distribution over a continuous feature space in early visual areas (Beck et al., 2008; Smith, 2016; Ratcliff, 2018). By effectively using such distributions for quantifying the degree to which trial-by-trial variations in brain signals are predictive of the corresponding judgements, we adapted the framework of so-called choice probabilities (Britten et al., 1996; Chicharro et al., 2021) to a feature-continuous task.

Our encoding models also allowed us to go beyond studying encoding itself and to explicitly assess the similarity between the stimulus-encoding and the report-encoding. For this we tested whether the information encoded by the report model when guessing could be used to infer the stimulus motion direction in a set of cross-prediction analyses (Figure 8). This procedure relies on the assumption that if the model estimated during one condition allows to predict data from another, then the brain activity patterns elicited by the two conditions are similar (Cichy et al., 2012; Kaplan et al., 2015; Levine & Schwarzbach, 2017). Note that for 100% and intermediate coherence levels there is a strong correspondence between the stimulus and corresponding reports (see e.g. Figure 2-B). This would make any correlation between stimulus and response encoding trivial. Thus, we limited our generalization analysis to the “guessing” condition, i.e. the reports for 0% coherence, which were not contaminated by stimulus-related signals. By using information encoded from this report model, we were able to predict the stimulus direction at intermediate and full coherence. We refer to this effect as model generalization. In contrast, we refer to model consistency as the ability of the report to predict reports in other coherence levels, but this is not a key focus here (see Figure 8). Our generalization analysis was successful, thus indicating that the sensory information and the choices are encoded in a similar representational feature space. This would suggest that the neural mechanisms recruited by visual areas to support perceptual decisions are, to some extent, the same used to encode visual stimuli. Whether this generalizes to discrete choice options or to other forms of report remains an open question.

Using our stimuli and task we did not find any significant motion direction or choice-related signal in area MT+. Previous studies have yielded somewhat inconsistent results on what to expect. There are numerous studies indicating the key role of MT+ in motion perception (Newsome & Parè, 1988; Rees et al., 2000; Braddick et al., 2001; Kamitani & Tong, 2006; Serences & Boynton, 2007; Sousa et al. 2021). However, other researchers employing multivariate pattern analysis and fMRI also did not find direction of motion information in MT+ (possibly due to short viewing times of stimuli, see e.g. Hebart et al., 2012, but see also Wang et al., 2014). Others have also found considerably lower levels of direction-related information in neuroimaging signals recorded from MT+ compared to V1 (Kamitani & Tong, 2006; Serences & Boynton, 2007). Please note that the relationship between single cell tuning and single voxel BOLD response profiles requires additional assumptions on how the voxel samples population of tuned sensory neurons (Nevado et al., 2004; Haynes, 2015; Sprague et al., 2018; Gardner & Liu, 2019, Sprague et al., 2019). For example, our negative finding could reflect the fact that the spatial distribution of cells encoding stimulus directions or choices in MT+ might not have been sufficiently anisotropic to differentially influence the fMRI signals. Alternatively, it could also reflect the continuous or graded nature of the task.

There are two other factors besides the continuous decisions that might help explain why choice-related information is only observed in early visual areas with our design. First, in our task specific motor preparation is impossible due to the employment of a visual comparison stimulus that rotated independently. While some studies have decorrelated choices and specific motor commands (e.g. Bennur & Gold, 2011; Bode et al., 2013) this is typically done by post-cueing a variable stimulus-response mapping and thus, the preparation of the response can be at least conditionally prepared (i.e. the motor command can be prepared in form of a differential response to a mapping cue).

A second difference is that the nature of our task involves a brief delay between the perceptual decision and the report in the absence of any possibility for motor preparation. Solving such a task might be achieved by briefly memorizing the target stimulus. In human neuroimaging studies of visual working memory sensory regions have been shown to encode stimulus features across short delays (Serences et al., 2009; Riggal & Postle, 2012; Pratte & Tong, 2014; Christophel et al., 2017), which is consistent with neuroimaging evidence from feature- continuous perceptual tasks involving a working memory component (van Bergen et al., 2015).

To summarize, our combination of a continuous feature task and fMRI encoding models suggested that early visual areas, but not MT+, allowed to reconstruct both continuous physical motion stimuli as well as continuous choices. Taken together, our results indicate that perceptual decisions regarding continuous sensory features might be encoded in early visual areas, potentially akin to visual working memory signals in sensory areas.

## MATERIALS AND METHODS

### Preregistration

The hypotheses, methods and analyses employed in this study were preregistered at https://osf.io/e2bvn before analyzing the data. Any additional exploratory analyses that go beyond what is specified in the preregistration are explicitly marked as such below.

### Data and Code availability

The data needed to reproduce the main analyses described in the paper are openly available on OSF at https://osf.io/vcmdg/ (DOI 10.17605/OSF.IO/VCMDG), whereas the code is available on Github https://github.com/RiccardoBarb/GPR_fMRI.

### Participants, payment and exclusion criteria

We recruited participants from several sources. Some were contacted using an internal mailing list consisting of people who previously participated in fMRI experiments in our lab. Others were recruited from Facebook groups for English-speaking jobs in Berlin and Berlin university students. All of the participants gave written informed consent, were paid 7€/h for the behavioral training session and 10€/h for the fMRI sessions. Those who completed all of the experimental sessions (1 training + 2 fMRI) received an additional bonus of 50€. The research protocol was conducted in accordance with the Declaration of Helsinki and approved by the local psychological ethics committee.

We selected healthy right-handed subjects with no history of neurological or psychiatric diseases. Furthermore, following our previous studies (Töpfer et al., 2022), we decided to exclude participants *prior to scanning* on the basis of their performance in a behavioral training session.

1. A participant was excluded if they were not sufficiently precise in the indication of motion direction, which was defined as the the 95% percentile of the Δ*x* distribution (see below, eq.1) in the full coherence condition exceeding a cutoff of 36.5°.
2. We also excluded participants that were not able to correctly perceive the stimulus in another, more systematic way. We had previously observed that some subjects frequently mistake a motion direction with its 180-degree opposite, a phenomenon that would be mistaken for a guess in most conventional categorical motion judgement tasks. For this reason, we employed a von Mises mixture model (vMMM) to quantify the frequency of reports of opposite direction (ROOD; Töpfer et al., 2022). ROOD rates exceeding 5% at full coherence led to the exclusion of a participant.

We initially collected behavioral training data from 41 subjects. Of these, 13 were excluded after the training phase prior to scanning following the abovementioned exclusion criteria. Three participants didn’t complete the MRI sessions for technical reasons and were excluded from the subsequent analyses; thus, 25 participants completed the fMRI sessions. One participant with low behavioral performance in the training session was accidentally included in scanning and was subsequently removed leading to a total of 24 participants who successfully concluded the experiment according to our pre-defined criteria (9 females; age range: 18-34; mean age: 25.6; SD: 4.6). We used the data of one participant to develop and check our Gaussian Process Regression (GPR) pipeline. In order to avoid any circularity or overfitting this participant was not included in the final analyses, which led to a total sample size of 23 subjects considered for all the statistical analyses.

### Stimuli: general features

The random dot kinematograms (RDKs – Figure 1) consisted of white dots moving inside a circular aperture on a black background. The aperture was centered on the screen and had an inner diameter of 2.5 dva and an outer diameter of 15 dva. Alpha blending was applied at the borders to avoid sharp contrast boundaries. For this, the luminance of the dots was progressively reduced before they wrapped around the other side of the annulus. The dot size was 0.1 dva (Braddick, 1973). The motion speed of the dots was 6°/s (van de Grind et al., 1983; Geisler, 1999) and the dot density was 1.6 dots/dva^2^ (Downing & Movshon, 1989), leading to a total of 275 dots. A white bullseye fixation target (Thaler et al., 2013) was placed in the central aperture spanning 0.25 dva. The mean luminance measured on the white center of the bullseye was 17.5 cd/m^2^. The mean luminance measured on the black background was 0.206 cd/m^2^.

### Stimuli: directions of motion

To pseudo-randomize the directions across trials, while maintaining the continuous nature of the task, we separated the stimuli into 8 hidden randomization bins. Each bin divided the stimulus space into equal portions of 45°. The bin edges were set at 337.5°, 22.5°, 67.5°, 112.5°, 157.5°, 202.5°, 247.5°, 292.5° (0° pointing up, 90° pointing right). Within a bin, the direction of motion was uniformly randomly distributed. In this way we made sure that the motion direction varied continuously across trials, while respecting some experimental constraints. We used an equal amount of trials for each subject in each directional bin, the same bin did not occur more than twice in a row and the same coherence level was not presented more than three times in a row.

### Training session: experimental setup

In order to train participants on the task lying-down in supine position (as during MR-scanning), we used a custom-built mock scanner. The training phase of the experiment took place in a dimly illuminated room (mean background luminance as measured on a white wall: 0.0998 cd/m^2^), where participants were lying in this mock scanner. They placed their head on a pillow and viewed a DELL LCD monitor 35 cm wide through a reflecting mirror. The monitor was set with 60 Hz refresh rate and a resolution of 1024x768 pixels. The stimuli were generated and presented using MATLAB R2016a (The MathWorks Inc.) and Psychtoolbox 3 (Brainard, 1997; Kleiner et al., 2007). For behavioral training, participants had their right hands on a standard computer keyboard placed on their hips.

### Training task

For the training session, each trial started with the presentation of a fixation bullseye which remained present throughout the whole duration of the trial (see Figure 1). Participants were instructed to fixate the center of the bullseye for the entire duration of the trial. After 0.5s, participants were presented for 2s with a random dot motion stimulus (RDK) that had a different direction of motion and coherence level for every trial. The direction of motion was continuously distributed between 0° and 360° and its order of presentation was subject to constraints (see above). There were five different coherence levels in the training phase: 0%, 12.5%, 25%, 50%, 100%. After termination of the stimulus, participants gave a judgement of motion direction using a report that employs a perceptual frame of reference with a visual comparsion stimulus (see Figure 1). In a previous study (Töpfer et al., 2022) we observed that this method of responding avoided systematic biases that are observed when using continuous reports involving trackballs that employ a motor frame of reference. Specifically, after offset of the motion stimulus a self-moving rotating bar was presented inside the aperture. Participants were asked to indicate the net motion direction of the dots by pressing the response button as soon as they believed the bar on the screen to match the direction of motion they perceived. The bar pointed from the center of the aperture to the outer border of the stimulus (like the arm of a clock), starting from a random position in every trial. The bar was 7.5 dva in length. It was randomly chosen to rotate clockwise or counterclockwise around the central fixation at a speed of 0.2 cycles/second, and it kept rotating after the response was given so that the total rotation time was always 7.5s. Participants were instructed to always respond as precisely as they could, even if they were unsure. On some trials (catch trials), a portion of the rotating bar changed contrast after the response indication. Participants were instructed to press the response button as fast as possible when they detected the contrast change. The purpose of such trials was to make sure that participants were paying attention to the bar rotation throughout the entire duration of the response period (even after they indicated their response), while maintaining fixation. In the training task, participants’ responses were followed by a uniform inter-trial interval (ITI) of 1 or 2s, after which a new trial started. Subjects performed 9 blocks of 40 trials during the training phase. An additional block was used to estimate the exact level of coherence that yielded intermediate performance using the QUEST staircase method (Watson & Pelli, 1983).

### fMRI experimental task

Participants who completed the training and matched our performance requirements (see section *Participants and exclusion criteria*) were scheduled for 2 different MRI sessions on 2 different days. Participants performed a total of 10 experimental runs (5 runs in each session). The structure of the training and the experimental tasks were essentially the same, except for the inter-trial interval (ITI), which was chosen between 3s, 5s, 7s and 9s, where ITI frequency was exponentially distributed (shorter ITIs were more likely than longer ones). Moreover, in the scanning sessions, the RDK was presented with three coherence levels instead of five: 0%, an intermediate coherence level estimated for each subject from the training phase data (mean coherence level: 19.47%; SD: 5.3%), and 100%. Each coherence level was presented 16 times in each run which resulted in 48 trials per run for a total of 10 runs. Participants were required to maintain fixation for the entire duration of the trial. Their eye position was monitored with an MRI compatible EyeLink1000+ eyetracking system.

### fMRI localizer task

After the experimental runs, participants performed 2 runs of an MT-localizer task on each day, for a total of 4 runs on both days. In the localizer, their task was to passively view the presented stimulus while maintaining fixation. After 0.5 s of bullseye, they viewed an 8s RDK at 100% coherence with random directions changing at a frequency of 2Hz, followed by 8 s RDK at 0% coherence. The ITI was implemented in the same way as in the main fMRI task. Participants were instructed to fixate the center of a bullseye for the entire block. The total number of blocks for each run was 32 (16 coherent, and 16 incoherent). This particular design has been proven effective in eliciting a strong BOLD signal in motion-sensitive visual areas (Braddick et al., 2001).

### MRI data acquisition

Functional MRI data were acquired on a 3T Siemens Prisma scanner (Siemens, Erlangen, Germany) equipped with a 64-channel head-coil, using a T2-weighted multi-band accelerated EPI sequence (from the Human Connectome Project - HCP) with a multiband factor of 8. The fMRI runs (TR = 800 ms, TE = 37 ms, flip angle = 52°, voxel size = 2 mm x 2 mm isotropic, 72 slices, 1.9 mm inter-slice gap) were preceded by a high-resolution T1-weighted MPRAGE structural scan (208 sagittal slices, TR = 2400 ms, TE = 2.22 ms, flip angle = 8°, voxel size = 0.8 mm^2^ isotropic, FOV = 256 mm). The MRI sessions took place over the course of two days. Each day comprised 5 experimental runs (805 whole-brain volumes per run) and 2 functional localizer runs (480 whole-brain volumes per run). The first 4 TRs were discarded to allow for magnetic saturation effects.

### Eye-Tracking data acquisition

Horizontal and vertical gaze position as well as the area of the pupil, were recorded from each subject’s dominant eye in the MRI scanner using an EyeLink 1000 + (SR-Research, sampling rate 1000 Hz) with long distance mount. Calibration took place before the experiment once at the beginning of every session.

### Data analysis

#### Behavior: Measures of performance accuracy

The absolute trial-by-trial circular response deviation from the target direction was used as a primary measure of performance accuracy. This is obtained by calculating

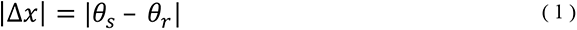

where *θ*_*s*_ is the stimulus direction and *θ*_*r*_ is the reported direction. Furthermore, as in previous studies we rescaled the absolute deviation in the range 0% - 100% (feature-continuous accuracy, FCA – see also Pilly & Seitz, 2009) following the formula:

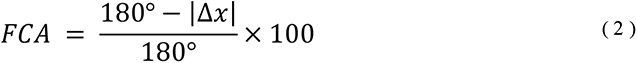

Here chance performance, i.e. randomly guessing the continuous direction, corresponds to an average FCA of 50% (or |Δ*x*| = 90°) and perfect performance, i.e. identically matching the presented direction, corresponds to an FCA of 100% (or |Δ*x*| = 0°). This approach has the advantage of providing a trial-by-trial measure of performance, which is interpretable at all coherence levels (including 0% coherence) and facilitates the comparison with more conventional 2-choice accuracies. An alternative to this approach would consist in fitting a mixture model to obtain an estimate of detection and guessing (Zhang & Luck, 2008; Bae & Luck; 2019; Töpfer et al., 2022). However, at 0% coherence, when there is no motion direction information available and subjects are purely guessing, the model fit would provide uninterpretable estimates for the detection parameter (Töpfer et al., 2022).

#### Localizer: Definition of ROIs

The fMRI data from the localizer runs were first spatially realigned, coregistered to individual anatomical images (Glasser et al., 2013) and spatially smoothed using a Gaussian kernel with an FWHM of 6mm. For each subject, we then modelled the activity during the localizer in each voxel using a general linear model implemented in SPM12. For each of the 4 runs, we included 1 regressor for coherent motion and 1 for incoherent motion, as well as 6 regressors of no interest to account for participants’ head movement. We then performed two univariate analyses: The first assessed in which voxels the BOLD signal was stronger during coherent compared to incoherent motion (for definition of MT+, see below). The second assessed where both coherent and incoherent motion activated voxels above baseline (for early visual and parietal brain regions, see below). The statistical maps obtained with this contrast were corrected for multiple comparison and thresholded at p < 0.05 (FWE). ROIs can be seen on the left part of Figure 6.

*Early visual cortex (EVC)*: Early visual cortex was defined based on a combination of a spatially normalized functional mask of motion-related activity and an anatomical mask defined by the union of V1, V2 and V3. Unlike MT+ masks, the functional EVC mask was defined at the group level because we didn’t expect significant inter-individual differences in the activation maps elicited by our contrast. Specifically, the functional activation elicited by both localizer stimuli (coherent and incoherent against baseline) constituted a large cluster (p < 0.05, FWE) at the occipital pole. Please note that this voxel selection is independent of motion coding information. For the anatomical mask we employed the SPM Anatomy Toolbox (Eichkoff et al., 2005) to define an anatomical mask spanning the occipical areas hOC1, hOC2 (Amunts et al., 2000), hOC3d (Rottschy et al., 2007) and hOC3v (Kujovic et al., 2013). Our early visual ROI is defined as the intersection between the functionally and the anatomically defined masks.

*Area MT+*: The motion complex MT+ was identified as the set of voxels activated more to the coherent than the incoherent localizer stimuli within a sphere (r = 10mm) located in the center of the significant clusters lateral to the parietal-occipital sulcus bilaterally (see Figure 6 for the MT+ ROI of an example subject).

*Parietal areas*: We used the SPM Anatomy Toolbox (Eichoff et al., 2005) to further select voxels from three different subregions of parietal cortex that have previously been reported as informative about behavioral choices in similar perceptual decision-making experiments (Bode et al., 2013; Hebart et al., 2012, 2016): Superior Parietal Cortex (SPC - areas 5L, 5M, 5Ci, 7A, 7PC, 7M, 7P; Scheperjans et al., 2008), Inferior Parietal Cortex (IPC - areas PFop, PFt, PF, PFm, PFFcm, PGa, PGp; Caspers et al., 2006) and Intraparietal Sulcus (IPS - areas hlP 1-3; Choi et al., 2006; Scheperjans et al., 2008).

#### Main experiment: Selection of trials based on eye-tracking

In order to avoid potential eye-movement confounds in our main fMRI analysis we checked that participants maintained fixation using the eye-tracker data. For this, in a first step we used a preprocessing pipeline adapted from Urai and colleagues (2017). Missing data and blinks were not interpolated for fixation control. The standard deviation of the gaze position was estimated for every run and every subject. We obtained the probability density function of the distribution of all standard deviations of eye positions, collapsed across all subjects and runs, using a kernel density estimation. A noise threshold was defined by estimating the inverse of the cumulative density at the probability of 0.9. This is equivalent to excluding the 10% of the noisiest runs based on the fixation analysis. Furthermore, trials where subjects exceed a deviation threshold of 2 dva for more than 200 ms during the stimulation period were rejected from the analysis of the neuroimaging data. The eye fixation control resulted in the exclusion of 295 trials, or an average of 2.3% of trials for each participant (mean number of trials: 11.13; SD: 14.51). Together with trials in which participants didn’t report their perceived direction, we excluded an average of 2.5% trials per subject (mean number of trials: 12.17; SD: 14.42). A repeated-measures ANOVA revealed no significant difference in the amount of excluded trials across coherence levels (F (2,44) = 1.296, p = 0.284).

#### Main experiment: Statistical analyses of behavioral performance

In order to evaluate the effect of coherence on behavioral performance, we performed a repeated-measure ANOVA on performance accuracy (FCA – see eq. 2), with coherence level as within-subject factor. The test was performed with JASP.

#### Main experiment: FMRI data analysis

The fMRI data analysis of the main experimental task was performed in MATLAB using SPM 12, the GPML toolbox (Rasmussen & Nickish, 2010) and custom functions. Before the analysis, data were motion corrected and coregistered to anatomical images. After this preprocessing, the analysis proceeded in four steps: (1) trial-wise and voxel-wise GLM; (2) voxel-wise Gaussian Process Regression (GPR) estimation; (3) searchlight-based stimulus and report reconstruction; (4) group level analyses.

*Trial-wise GLM:* For each participant, we modeled the fMRI signal acquired in each voxel during the 2s of the stimulus period with a trial-wise GLM (Rissman et al., 2004). Our model consisted of one regressor per trial and 6 head-motion regressors of no interest for each run.

*Voxel-wise GPR estimation:* In a next step we computed what we refer to as Full Distribution Tuning Functions (FDTF) for each voxel to assess how the estimated trial-wise responses in that voxel were modulated either by the stimulus direction *θ*_*s*_ or by the reported direction *θ*_*r*_. The key idea of the FDTF over and above conventional voxel encoding models is to not only estimate the mean response in a given voxel as a function of direction, but to estimate the entire ensemble of direction-conditional likelihood functions (for details see below). For this we used Gaussian Process Regression (Rasmussen & Williams, 2005). The trial-wise parameter estimates were entered into two separate cyclic GPR, models, one for the corresponding stimulus motion direction *θ*_*s*_ and one for the reported direction *θ*_*r*_. This yielded a feature-continuous model of the entire distribution of fMRI responses for each direction, akin to a voxel-tuning function that includes not only the mean but also the distribution at each direction. The GPR has the advantage that it does not pre-suppose a fixed number of sensory channels per voxel (Brouwer & Heeger, 2009; van Bergen et al., 2015). This procedure was repeated for each coherence level, leading to the estimation of two separate sets of FDTFs for each voxel, three coherence levels and physical vs reported directions (six estimated models in total).

To be more specific, consider the t = 160 total trials for one coherence level across runs. Then, let *β̂_j_* be the *t* × 1 vector of trial-wise parameter estimates in a single voxel *j* and let *θ* be the corresponding vector of stimulus or reported motion directions:

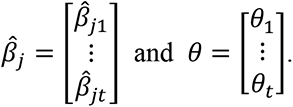

We assumed that the response amplitude of each voxel *j* during each trial *i* was a function of this trial’s direction of motion *θ*_*i*_ ∈ Θ = (0,2*π*] and the voxel-specific kernel parameters *L*_*j*_, plus the normally distributed noise term *ε*_*ji*_ :

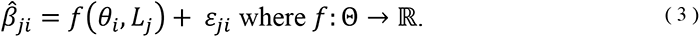

The function *f* is intentionally specified unconstrained, because it was estimated using voxel-wise GPR of responses *β̂*_j_ against directions *θ*, such that each voxel *j* obtained a unique response profile *f*_*j*_. The estimation of such voxel-wise response profiles was performed separately for each coherence level. For this we only used trials in which participants maintained fixation, as evaluated by the analysis of the eye-tracking data (see *Main Experiment: Selection of trials based on eye-tracking*). In order to avoid overfitting in the subsequent phases of the analysis (i.e. the searchlight reconstruction), we estimated the response profiles *f*_*j*_ by including the trials from all runs except one, and repeated the procedure until the trials from all runs were used for the model estimation (leave-one-run-out cross- validation scheme). Each iteration was based on a maximum of 9⁄10 × 160 = 144 datapoints (which could be maximally achieved when no data were excluded after fixation control).

*Searchlight-based stimulus and report reconstruction:* Next, we used the estimated tuning functions to reconstruct the direction for a set of independent test trials. We then combined the estimated *β̂_j_* parameters from a group of voxels within a searchlight (r = 4 voxels) into the matrix B̂, to predict the stimulus direction *θ*_*s*_ or the reported direction *θ*_*r*_ using a run-wise cross-validation procedure. The estimated voxel-wise response profiles *f*_*j*_ = *f*(*θ*_*i*_, *L*_*j*_) can be used to predict the 1 × *υ* vector of response amplitudes across voxels *j* in one trial *i*, i.e. one row of B̂:

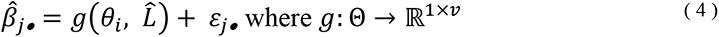

as well as the whole *t* × *υ* matrix of trial-wise parameter estimates B̂:

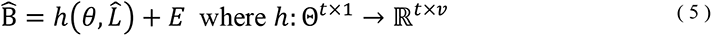

with *L̂* = {*L*_1_, …, *L*_*υ*_}, where *g* and *h* can be written in terms of *f* as:

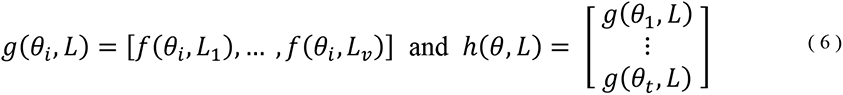

Let’s partition the trial-wise parameter matrix B̂ into training data B̂_train_ and test data B̂_test_ and let *L̂* = {*L*_1_, …, *L*_*υ*_}, be the set of estimated kernel parameters obtained from B̂_train_. Then, the residuals of the GPR model in the voxel-wise GPR equation are:

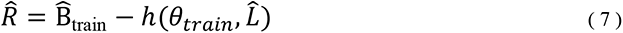

and estimated covariance is:

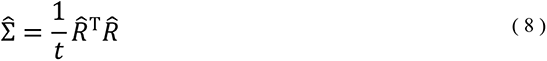

In the case that the number of voxels in a searchlight *υ* is larger than the number of trials across runs *t*, the matrix is not invertible, such that it has to be regularized. Here, we chose a shrinkage estimator by mixing in a diagonal matrix of sample variances

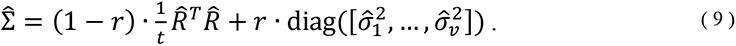

Where the mixing coefficient *r* was a function of the voxel-to-trial ratio:

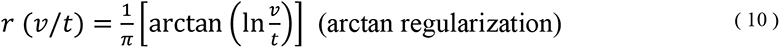

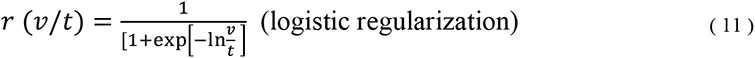

such that 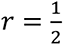 when 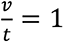 and 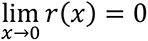 as well as 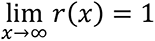.

Let’s consider the responses of all voxels in just one trial. According to the GPR model, those single-trial across-voxel responses are distributed as

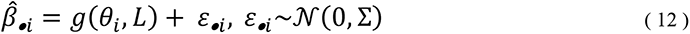

Which implies a multivariate normal log-likelihood function

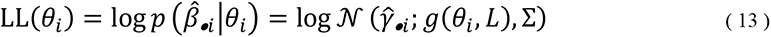

where 𝒩 (*x*; μ, Σ) is the probability density function of the multivariate normal distribution and Σ is the unknown *υ* × *υ* spatial covariance matrix.

The stimulus or the reported motion direction in a particular trial of the test set can be reconstructed by maximum-likelihood estimation (MLE), i.e. by simply maximizing the out-of-sample likelihood function, given the in-sample parameter estimates *L̂* and Σ̂:

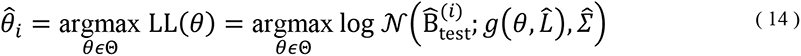

where 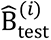 is the *i*-th row of B̂_*test*_, containing parameter estimates from the *i*-th test trial. To efficiently determine *θ̂*_*i*_, we perform a grid-based search, ranging *θ* from 1° to 360° in steps of 1° and evaluating LL(*θ*) at the corresponding values. This approach is similar to inverting a set of forward encoding models (Thirion et al., 2006; Brouwer & Heeger, 2009; Naselaris et al., 2011; Haynes, 2015; Sprague et al., 2018; Kriegeskorte & Diedrichsen, 2019).

*Reconstruction performance evaluation:* The outcome of the reconstruction is the matrix of predicted directions *θ̂*]

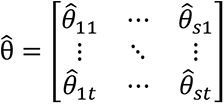

where *t* is the number of trials and *s* is the number of searchlights.

Following our preregistration protocol, we evaluated the reconstruction performance in terms of feature-continuous accuracy (FCA, eq. 2). For each searchlight and trial, we compared the true (stimulus or report) direction *θ*_*t*_ and the predicted direction *θ̂*_*t*_ in terms of absolute angular deviation, rescaled into the range 0-100%, according to equations (1) and (2). We then computed the averaged FCA across trials. This measure works well with balanced independent variable distributions, such as that of the stimulus motion directions. However, in order to avoid spurious above-chance reconstruction performance in case of an unbalanced distribution of the dependent variable (i.e. during the reconstruction of participants’ reports which are not equally distribution across the directions, especially at lower coherence levels), we computed a balanced version of FCA (BFCA):

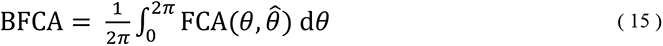

where the integral is calculated using trapezoidal numerical integration across the sorted directions of motion *θ* and reconstructions *θ̂*. Note that, in case of balanced labels, the use of FCA or BFCA produce virtually identical results (See Appendix 2 for details).

#### Statistical analyses: ROIs

In order to evaluate how the stimulus and the report reconstruction performance were affected by coherence in early visual cortex, MT+ and parietal areas, we evaluated the effect of coherence level on the reconstruction performance for five bilateral ROIs (see *Localizer: Definition of ROIs*). Parietal areas encode a variety of task-related variables (Bennur & Gold, 2011; Rigotti et al., 2013; Park et al., 2014; Brincat et al., 2018), including perceptual choices in previous studies where decisions only had few alternative options (Liu & Pleskac, 2011; Hebart et al., 2012, 2016). We expected an effect of coherence on report reconstruction in visual areas and MT+ reflecting the increased correlation between stimulus and response for higher coherence levels. To test these hypotheses, we first computed the average reconstruction performance for each label (stimulus and report) in each ROI for each subject. We then submitted the data to five independent repeated-measures ANOVAs (one for each ROI) with coherence level and reconstructed label as repeated measure factors. The statistical tests for the ROI analyses were performed with JASP (JASP Team, 2020).

#### Additional exploratory analyses: fMRI data

In order to deepen our understanding of the relationship between coherence level and reconstruction performance, and to explore the similarity between the stimulus and report GPR models, we further performed two exploratory analyses. The analyses were performed on an ROI defined by the intersection of voxels with an average above-chance reconstruction performance for both the stimulus and the report (thresholded at p < 0.001, uncorrected).

- *Interaction between coherence level and reconstructed labels.* We wanted to clarify whether the coherence level affected the stimulus and the report reconstruction in the voxels that code for both. For this, we computed the mean reconstruction performance of the ROI for each coherence level and each reconstructed label. Finally, we performed a repeated-measures ANOVA to test for an interaction effect between coherence level and reconstructed label.
- *Model generalization.* To further explore the similarity between the encoding of stimulus and report information in the brain, we performed a series of cross-prediction analyses. The principle of this analyses is the same as in other examples of cross-classification (Cichy et al., 2012; Kaplan et al., 2015; Levine & Schwarzbach, 2017). It involves the use of the GPR estimated in one condition (e.g. the report model estimated with the data acquired in the 0% coherence condition) to predict the data of a different one (e.g. the stimulus directions at 100% coherence condition). If the model generalizes, then there is evidence that the pattern of brain activity is similar across the two conditions. For each subject we tested how well the report model estimated in the 0% coherence condition (which is independent of the physical stimulus) would allow to predict the stimulus identity in each coherence level and vice versa (averaged). The procedure was similar to the one described in *Main experiment: fMRI data analysis - Voxel-wise GPR estimation* and *Searchlight based stimulus and report reconstruction*. However, we ran the analysis on a limited number of voxels (see above). Since we were primarily interested in the generalization of the report model estimated at 0% coherence to the stimulus model estimated at 100% coherence (see Discussion), we tested whether the generalization performance of this pair of conditions were above chance by computing a one tailed t-test.

## APPENDIX 1

### Statistical analyses: searchlight-based reconstructions and effect of coherence

We also performed additional whole-brain searchlight decoding to identify regions potentially outside our pre-defined ROIs that might encode stimulus-related or choice-related information. For this, the single-subject BFCA maps resulting from the searchlight reconstructions were smoothed using a Gaussian kernel with a FWHM of 6 mm and spatially normalized with SPM12. In order to identify searchlights in which the reconstruction performance was significantly above chance (i.e. BFCA > 50%), we used two one-way factorial designs (one for stimulus and one for report) with coherence level as a within-subject factor. For each model, we specified three t-contrasts. In this way we could identify clusters with significant above-chance information for each coherence level. We expected the reconstruction performance of searchlights located in visual areas, to decrease as a function of decreasing coherence. We also expected to identify clusters of voxels carrying information about perceptual judgements in visual (Britten et al., 1996; Serences & Boynton, 2007; Hebart et al., 2012; Sousa et al., 2021) and parietal areas (Gold & Shadlen, 2007; Brincat et al., 2018; Hebart et al., 2012, 2016; Levine & Schwarzbach, 2017). We further predicted the coherence level to have an effect on report reconstruction performance as well. We evaluated the effect of coherence on the reconstruction performance by inclusively masking the voxels that showed an average effect of reconstruction across coherence levels (p < 0.001, uncorrected). This procedure was done separately for stimulus reconstruction and report reconstruction. Please note that this analysis is not circular because the test for an effect of coherence is orthogonal to the test for average reconstruction performance.

The whole-brain searchlight reconstruction at 100% coherence revealed stimulus information from voxel clusters located in the left (FWEc, p < 0.05, K = 1677; cluster-defining threshold p < 0.001) and in the right occipital cortex (FWEc, p < 0.05, K = 1024; cluster-defining threshold p<0.001). For the intermediate coherence condition and for the 0% coherence condition, we found no searchlights that were significantly predictive of the stimulus motion direction. Note that in the 0% coherence condition, the stimulus has no global motion direction, thus a chance-level reconstruction performance is to be expected. In the left and right occipital cortex, we also found clusters of voxels informative about participants’ reports for the 100% coherence condition (left: FWEc, p < 0.05, K = 1049; cluster-defining threshold p < 0.001; right: FWEc, p < 0.05, K = 864; cluster-defining threshold p < 0.001) as well as for the intermediate coherence condition (left: FWEc, p < 0.05, K = 201; cluster-defining threshold p<0.001; right: FWEc, p < 0.05, K = 487; cluster-defining threshold p<0.001). For the 0% coherence condition, we were not able to identify clusters informative about participants’ reports.

**Appendix 1-Figure 1:**
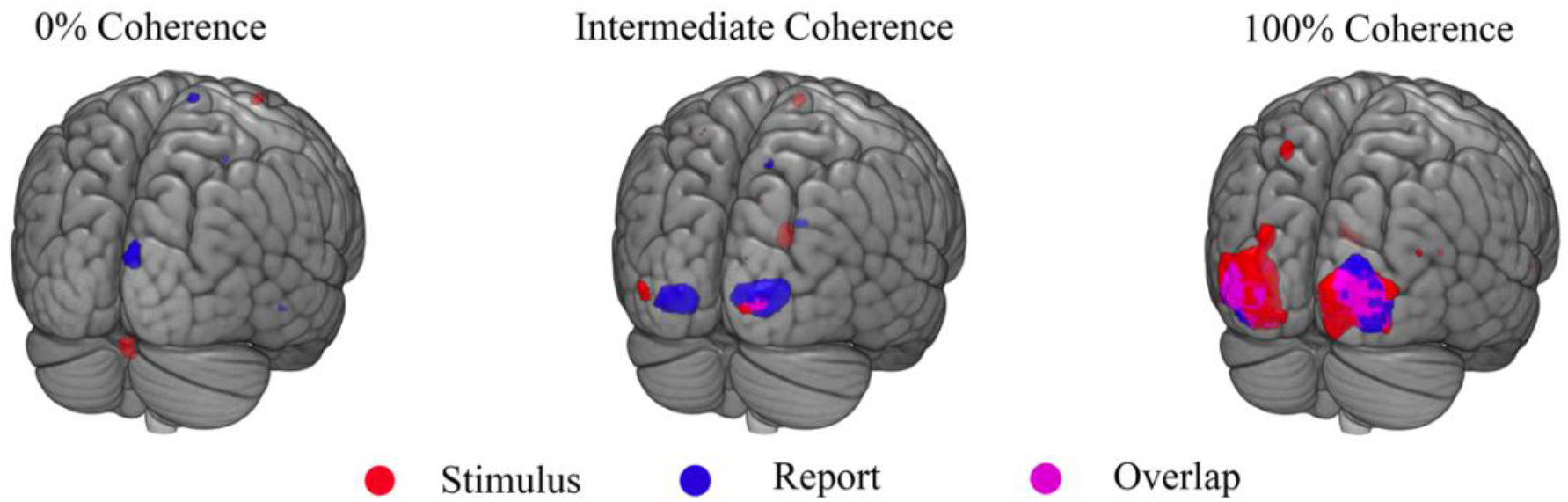
Searchlight-based accuracy maps (plotted here for BFCA, see Methods). The images show results of the searchlight-based stimulus and report reconstructions for three coherence levels (left: 0%; middle: intermediate; right: 100%). The searchlights are mapped with different colors: red indicates significantly above-chance reconstruction performance for stimulus, blue indicate above-chance reconstruction performance for report, and purple indicate the overlap between the two. Please note that the maps are shown for display purposes (for visualization thresholded at p < 0.001 and not corrected for multiple comparisons).

Since neurons in early and extrastriate visual areas are tuned to motion directions (Albright et al. 1984; Movshon & Newsome, 1996; Nichols & Newsome, 2002), we reasoned that if population-level measurements of neural activity obtained from single voxels reflects this property (Nevado et al., 2004; Haynes, 2015; Sprague et al., 2018), the stimulus reconstruction performance should be maximum in the 100% coherence condition, and progressively decrease at intermediate coherence. At 0% coherence instead, the stimulus has no net motion direction and the reconstruction performance should be at chance level. We therefore expected to identify a main effect of coherence on the stimulus reconstruction performance in visual areas.

**Appendix 1 – Figure 2:**
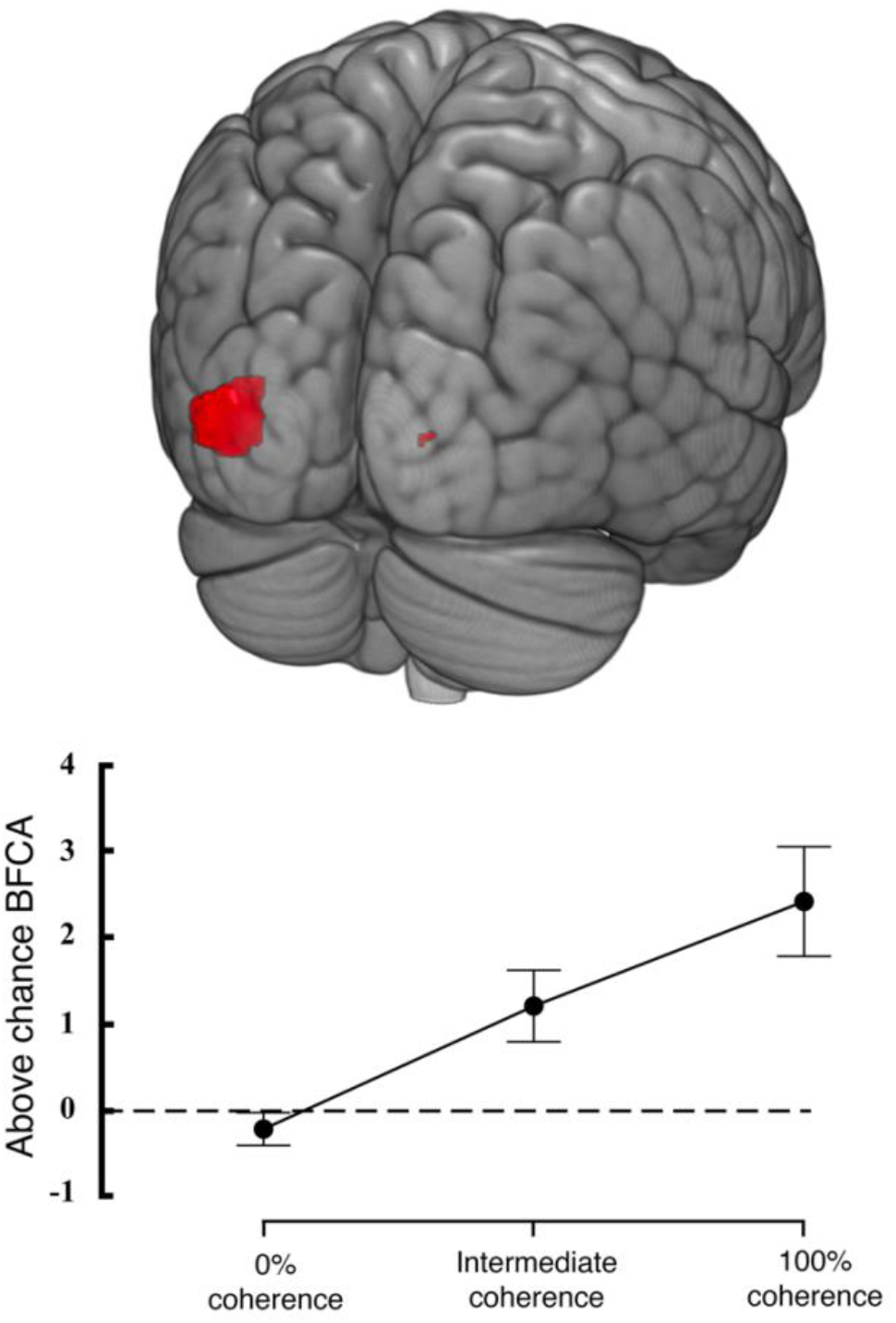
Effect of coherence on searchlight-based stimulus reconstruction performance. The picture on top shows clusters of voxels where coherence has a significant effect on stimulus reconstruction performance. The map is thresholded at p < 0.001, uncorrected for multiple comparisons. The plot on the bottom displays the averaged above-chance accuracy (BFCA minus baseline of 50%) extracted from the searchlights in which coherence had a significant effect on stimulus reconstruction performance, error bars are standard errors (N=23).

We identified a cluster of voxels located in the left occipital pole, where coherence level had an effect on the performance in stimulus reconstruction (FWEc, p < 0.05, K = 262; cluster- defining threshold p<0.001). Similarly, we predicted that a possible effect of coherence on the report reconstruction might be present, and driven by the expected correlation between the stimulus identity and participants’ report, when the stimulus is clearly visible. However, we did not find clusters that showed such effect for the report reconstruction performance.

### Control analysis: stimulus and report reconstruction from eye-tracking data

The use of motion stimuli such as our RDK might trigger involuntary eye movement (Cohen et al., 1977) that can be informative about the direction of perceived motion (Wilbertz et al., 2018). Eye movements also have an effect on brain activity measured with fMRI and can thus constitute a potential confound (Merriam et al., 2013), even when participants are specifically instructed to maintain fixation (Thielen et al., 2019). For this reason, we tested whether the recorded gaze position (x and y ordinates), was informative about the physical stimulus direction or participants’ perceived direction. For this analysis we used the gaze position of 21 out of 23 subjects who participated in the main fMRI experiment (we couldn’t record the traces of two participants for technical reasons). We reasoned that if the gaze position is systematically correlated with the presented motion direction or with participants’ reports, the x and y ordinates should exhibit a specific position profiles, that can in turn be used to perform stimulus and report reconstruction. In order to check whether the pattern of participants’ eye movements was related with the stimulus or the reported motion directions, we estimated such profiles with a cyclic version of the GPR (see *Main experiment: fMRI data analysis* - *Voxel-wise GPR estimation* in the Materials and Methods section) and performed stimulus and report reconstruction at each time point with a procedure similar to the one adopted for the main fMRI analysis (see Materials and Methods: *Searchlight-based stimulus and report reconstruction*).

The preprocessing pipeline employed for this analysis was different from the one performed for fixation control (see *Main experiment: Selection of trials based on eye-tracking* in the Materials and Methods section). Blinks, detected by the provided software from Eye Link, were linearly interpolated using the approach described in Urai et al. (2017). The resulting traces were filtered for electronic noise using a Butterworth filter (low cut off 5Hz, high cut off 100Hz) following Thielen et al. (2018). The complete trace of each session was linearly detrended to account for drifts that appear due to the long continuous recordings during each session (each approximately 1.5h). An additional linear detrending was performed separately on each run, to counterbalance slow drifts in head position in the scanner. Periods of interest (500 ms before stimulus onset together with 2000 ms stimulus period) were combined across the two recording sessions. Data from the period of interest were baseline corrected using the pre-stimulus interval (500 ms up to stimulus onset).

After preprocessing, the x and y gaze positions of each subject were grouped by coherence levels (0%, medium, 100% coherence) resulting in a maximum of 160 trials per condition. To reduce computational time, we only considered time points during the stimulus period (2000 ms) and resampled the signal at 50Hz. For each time point we estimated two position-related profiles (one for each ordinate) by entering the trial-wise position value together with the corresponding stimulus motion direction *θ*_*s*_ or the reported direction *θ*_*r*_ into a cyclic version of the GPR. Please note that this procedure is very similar to the one previously described for the estimation of voxel-wise response profiles (see eq. 3 in the Materials and Methods section - where the parameter *β̂*_*j*_ represents now the trial-wise recorded position of each ordinate in a single time point). The estimation of the position profiles was performed with a leave-one-run-out cross-validation scheme, by only using trials in which participants were maintaining fixation (see *Main experiment: Selection of trials based on eye-tracking* in the Materials and Methods section).

The stimulus and report reconstructions were estimated using the same procedure described in the section *Main experiment: fMRI data analysis* - *Searchlight based stimulus and report reconstruction* in the Materials and Methods section. However, the estimated gaze position profiles instead of voxel response profiles, were used to predict the stimulus direction *θ*_*s*_ or the reported direction *θ*_*r*_ in a run-wise cross-validation procedure. Please note, that in this case it is not necessary to adopt regularization for estimation of the covariance matrix because the number of position profiles (one for x and one for y ordinates) does not exceed the number of trials across runs (see eq. 9 in the Materials and Methods section). The results were evaluated by testing if the averaged BFCA (see *Main experiment: fMRI data analysis - Reconstruction performance evaluation* in the Materials and Methods section) across subjects was above chance for each timepoint. Statistical analyses were corrected for multiple comparison by performing a cluster-based permutation test (Maris & Oostenveld, 2017).

The group-level average reconstruction performance for the stimulus and the report labels are depicted in Appendix 1- Figure3. We were not able to identify stimulus-related information in any of the three coherence levels. Instead, the evaluation of the report model indicates that the pattern of eye movements was informative about participants’ report in the 0% coherence condition. More precisely we were able to identify clusters of above-chance reconstruction performance, peaking after 1000 ms. Eye movement were not predictive of participant’s choices for the intermediate or 100% coherence levels.

**Appendix 1 - Figure 3:**
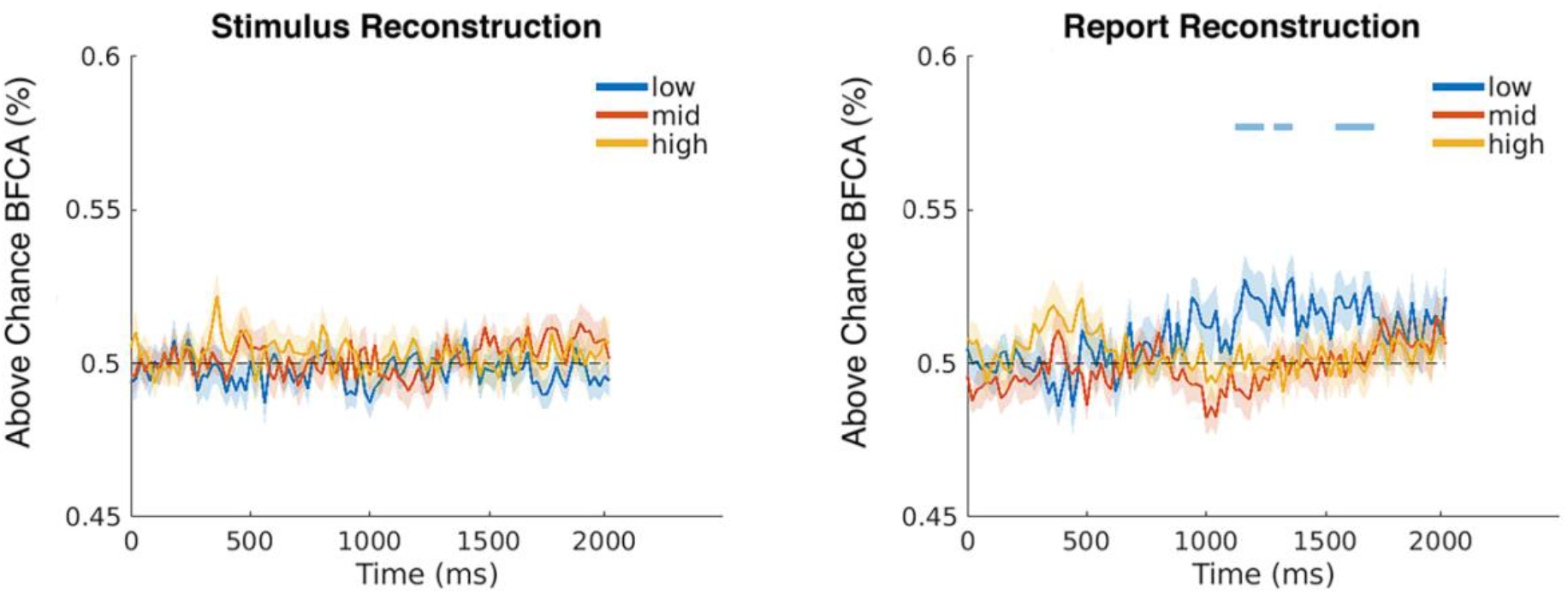
Group average (N=21) accuracy (expressed as BFCA, see Methods) at each time point relative to stimulus onset. The upper picture displays the stimulus reconstruction performance, the lower picture shows the report reconstruction performance for three coherence levels (blue: 0%; red: intermediate; orange: 100%). Shading around the individual curves indicates ±1 SEM. The light blue lines on top of the curves depict clusters of time points for which the reconstruction performance was greater than chance after correction for multiple comparisons. Please note that GPR estimated from eye-movements are predictive of the reports for the 0% coherence condition but not of those given at intermediate and 100% coherence. Such result together with those of our model consistency and model generalization analyses (Figure 8), suggest that eye movements were unrelated with the brain signals used to reconstruct participants’ choices in the 0% coherence condition (Figure 6).

## APPENDIX 2

### The need for BFCA

In order to evaluate the performance of our GPR-based reconstructions, we implemented a balanced version of FCA (BFCA – see eq. 15 in the Materials and Methods section). Our goal was to obtain a measure of performance that could be intuitively compared to a standard accuracy measure with values distributed between 0% and 100%. FCA is derived by rescaling the continuous values of the absolute angular deviation (see eq. 1 and 2 in the Materials and Methods section); see also Pilly & Seitz, 2009), to evaluate the reconstruction performance. The need for a *balanced* version of FCA was due to participants’ responses in the 0% coherence condition being unbalanced (Appendix 2 – Figure 1) as is often the case for reports, even despite the use of our sensory matching approach that minimizes such biases (Töpfer et al. 2022). In case of a standard classification analysis, training and testing a classifier with unbalanced labels make accuracy an unreliable measure of performance (Japkowicz & Stephen, 2002). More specifically, when the performance of a classifier is tested on an imbalanced dataset, it might lead to the misleading finding of significant above-chance performance of the classifier (Brodersen et al., 2010), simply because the classifier tends to reproduce the distribution of the training dataset.

**Appendix 2 – Figure 1:**
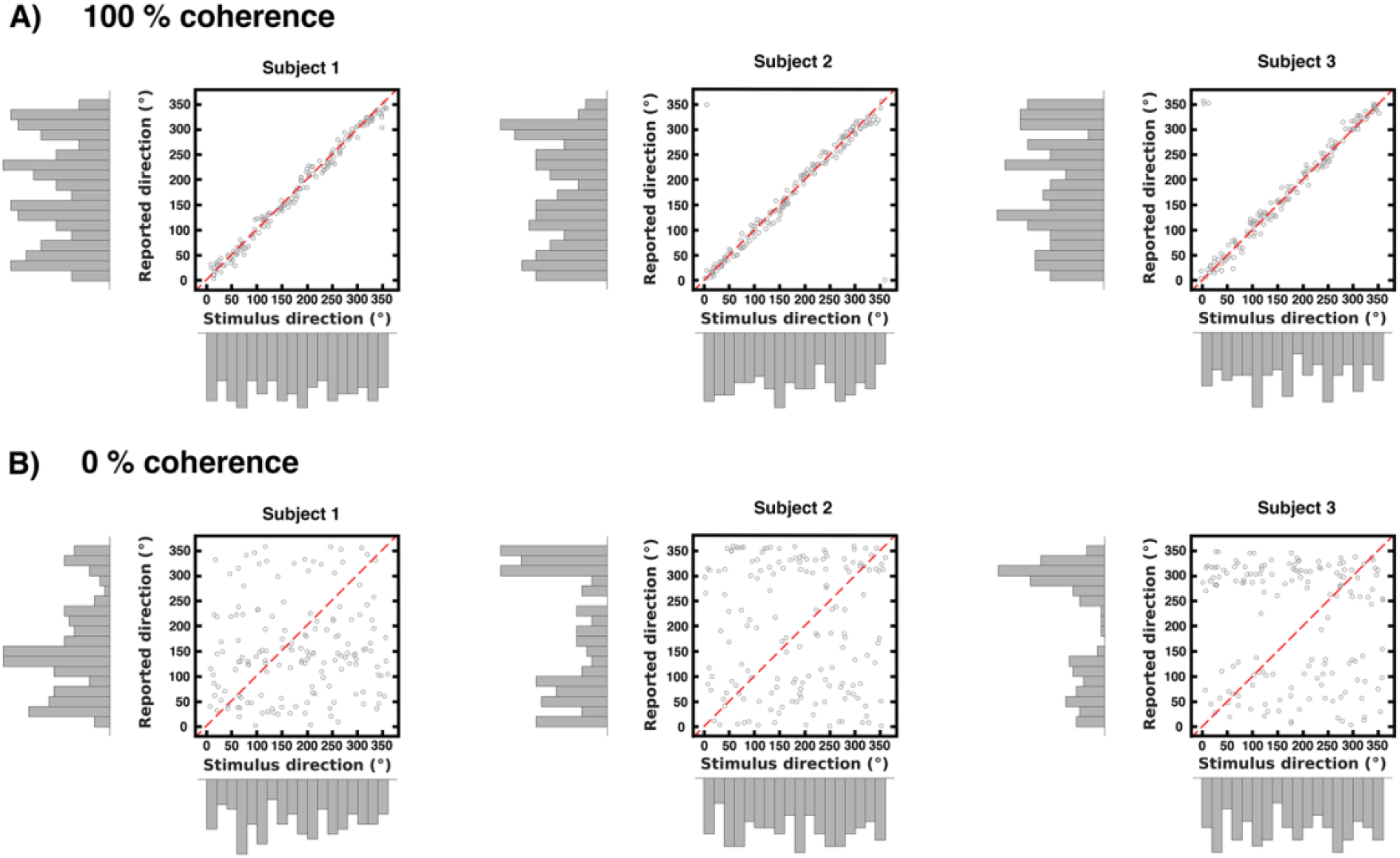
The scatterplots display the trial-wise reported direction of 3 example participants against the trial-wise motion direction, with the corresponding marginal distributions. A) Each plot on the top row shows the data distribution obtained from 160 trials in the 100% coherence condition. B) The bottom row shows the data distribution for the 0% coherence condition. Note that in this case the motion direction labels were generated following the randomization scheme described in the manuscript (section *Stimuli: directions of motion* in the Materials and Methods section), as no real motion direction was present in the stimulus.

### Comparing FCA and BFCA: simulation

In order to illustrate how BFCA and FCA are related with each other, as well as with the underlying independent variable distribution, we here show a simulation performed on synthetic data. In order to match the features of our experimental design, we simulated a *t* × 1 vector of trial-wise parameter estimates *β̂_j_* for a total of 1000 voxels where *t* = 160 total trials were generated across 10 runs. We also generated a vector *θ* corresponding to the independent variable (the stimulus or the report direction):

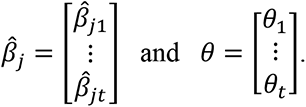

For the current simulation we distinguished four alternative scenarios:

1. ***θ*** **modulates** ***β̂***_***j***_**and the distribution of** ***θ*** **is balanced**. This situation corresponds to the hypothesized behavior of voxels sensitive to motion directions (as the stimulus directions are balanced across runs in our experimental design);
2. ***θ*** **modulates** ***β̂***_***j***_**and the distribution of** ***θ*** **is unbalanced**. This scenario corresponds to the hypothesized behavior of voxels sensitive to participants’ reports in our experimental design (as participants’ reports are unbalanced, especially at 0% coherence level);
3. ***θ*** **does not modulate** ***β̂***_***j***_**and the distribution of** ***θ*** **is balanced**. This should be the case for voxels insensitive to the stimulus direction. Such voxels should not produce spurious above-chance FCA when combined for searchlight-based reconstruction;
4. ***θ*** **does not modulate** ***β̂***_***j***_**and the distribution of** ***θ*** **is unbalanced**. We assume that this scenario could possibly produce spurious above-chance FCA when the voxels are combined for searchlight-based reconstruction.

We applied the same analyses described in the manuscript (see *Main experiment: fMRI data analysis* in the Materials and Methods section) to estimate voxel-wise response profiles using GPR and to perform the searchlight-based reconstruction using MLE. The simulated searchlights consisted of 241 voxels. We finally evaluated the reconstruction performance by using averaged FCA and BFCA.

The results of the simulation are summarized in Appendix 2 – Figure 2. We obtained an above-chance reconstruction performance for cases 1 (mean FCA: 92.31%, SD: 1.57; mean BFCA: 91.73%, SD: 1.9) and 2 (mean FCA: 92.72%, SD: 1.09; mean BFCA: 90.99%, SD: 1.91).

Interestingly, for case 3 the mean reconstruction performance is around chance for both measures (mean FCA: 50.36%, SD: 2.61; mean BFCA: 49.21%, SD: 2.56) whereas for case 4 the distribution of FCA is skewed toward right (mean FCA: 56.23%, SD: 4.25) whereas BFCA values are not (mean BFCA: 46.98%, SD: 6.06), as confirmed by two one-sample right tailed t-tests evaluating whether the mean values were greater than 50% (FCA > 50% : t = 46.278; p < 0.001; BFCA > 50%: t = -15.747; p = 1).

**Appendix 2 - Figure 2:**
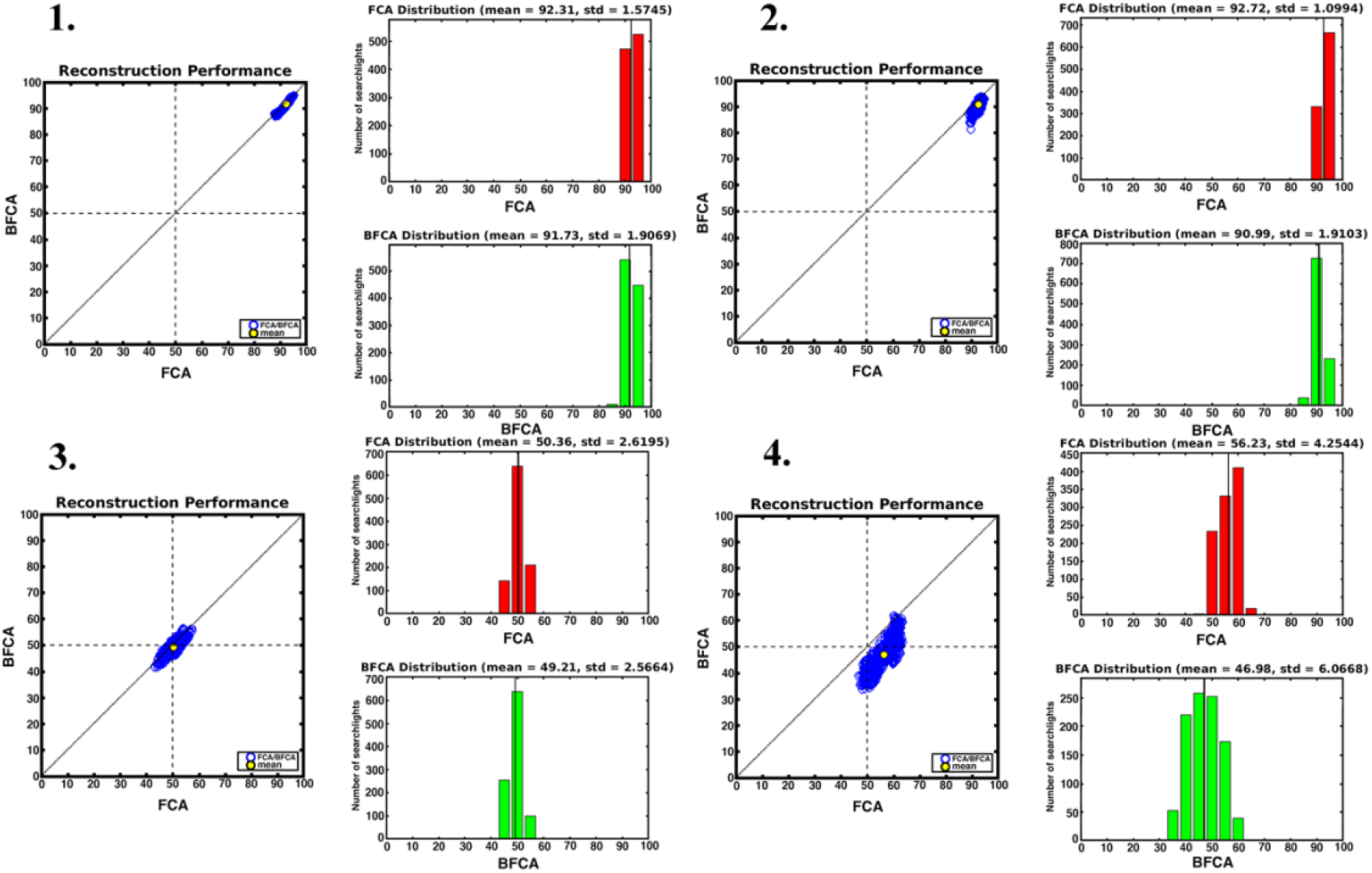
Comparison of reconstruction performances obtained with simulated data. The picture illustrates the distribution of FCA (red) and BFCA (green) in the four scenarios examined in the simulation. 1.

The relationship between FCA and BFCA for the condition in which *θ* modulates *β̂_j_* and the distribution of *θ* is balanced. 2. Condition in which *θ* modulates *β̂_j_* and the distribution of *θ* is unbalanced. 3. *θ* does not modulate and the distribution of *θ* is balanced. 4. *θ* does not modulate *β̂_j_* and the distribution of *θ* is unbalanced.

### Comparing FCA and BFCA: real data

We computed the result the whole-brain searchlight analysis for all of the 23 subjects both with FCA and BFCA as measures of reconstruction performance. The maps were obtained following the procedure described in *Main experiment: FMRI data analysis* in the Materials and Methods section. For the purpose of this analysis we only considered three main conditions:

1. **Stimulus labels at 100% coherence**. In this condition, the stimulus motion directions are balanced, therefore the reconstruction performance should be above chance only for the searchlights with voxels sensitive to motion directions. Because the distribution of the stimulus directions is balanced, we expect no difference between the reconstruction performance computed with FCA and BFCA.
2. **Stimulus labels at 0% coherence**. In this condition, the stimulus had no real motion direction, but each trial was assigned a motion direction, generated according to our randomization scheme (see section *Stimuli: directions of motion* in the Materials and Methods section) This results in a balanced label distribution. Because of this, no searchlight should result in above-chance reconstruction performance. Following the outcome of the simulation described above, we expect no difference between the reconstruction performance computed with FCA and BFCA.
3. **Report labels at 0% coherence**. Here, the labels assigned to each trial correspond to the motion directions reported by participants. Therefore, the distribution of the reports across trials reflects the idiosyncratic biases of each subject (see Appendix 2 – Figure 1 for an example). In this condition, because some participants’ choices lead to an unbalanced distribution of reported motion directions, we suspect that FCA leads to spurious above- chance reconstruction performance. Based on the outcome of the simulation, we hypothesize a difference in the reconstruction performance computed with FCA and BFCA.

We then used SPM12 to compare the FCA and the BFCA maps of the 23 participants. We performed three second-level paired t-tests to evaluate our hypotheses.

**Appendix 2 – Figure 3:**
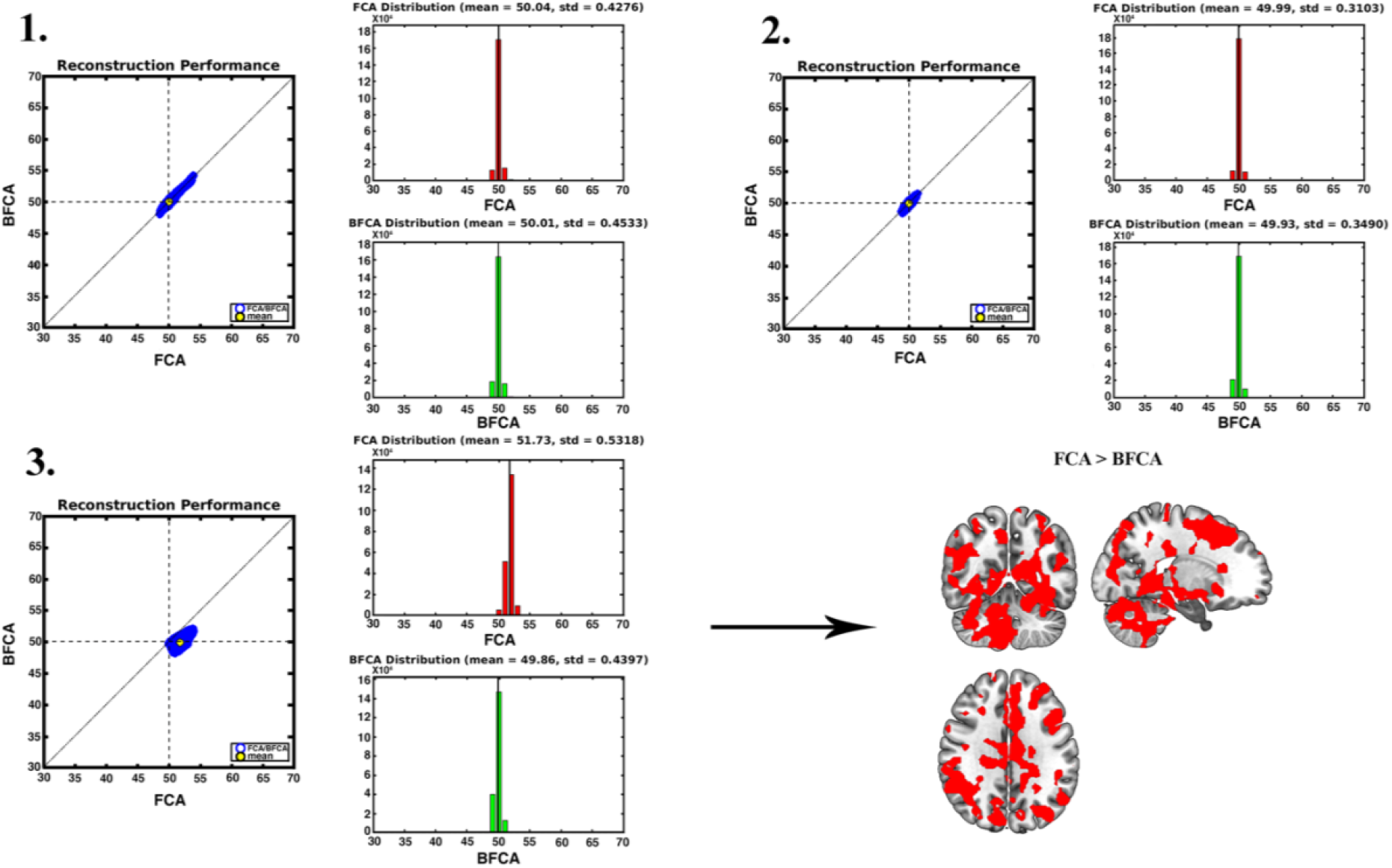
The plots illustrate the difference between reconstruction performances obtained with two accuracy measures, FCA and BFCA. 1-3 show the relationship between FCA and BFCA for the 100% coherence stimulus reconstruction, for the 0% coherence stimulus reconstruction, and for the 0% coherence report reconstruction respectively. Results are plotted for each searchlight, averaged across subjects (N=23). The brain map on the bottom right is obtained from a 2^nd^ level t-test evaluating searchlights where FCA was higher than BFCA in the 0% coherence report reconstruction (N=23). The map is thresholded at p<0.001, uncorrected for multiple comparisons.

The two measures (FCA and BFCA) were not significantly different when using the stimulus labels at 100% coherence and 0% coherence (Appendix 2 - Figure 3). However, the FCA was significantly different from BFCA when using the labels of the 0% reports in a large cluster covering various portions of the brain (FWEc, p < 0.05, K = 667990; cluster-defining voxel threshold p < 0.001). The results remained consistent even when we lowered the cluster- defining threshold to p < 0.0001 or p < 0.00001, with many significant clusters of smaller size scattered throughout the brain reaching significance level (FWEc, p < 0.05, smallest K = 70; cluster-defining voxel threshold p < 0.0001; FWEc, p < 0.05, smallest K = 20; cluster defining voxel threshold p < 0.00001).

## Acknowledgments

Funded by the Excellence Initiative of the German Federal Ministry of Education (Excellence Cluster Science of Intelligence), the BMBF (through the Max Planck School of Cognition), the DFG (GRK 2386 “Extrospection”), and a joint grant by the John Templeton Foundation and the Fetzer Institute. We would like to thank Philipp Sterzer and Tobias Donner for their valuable input.

## REFERENCES

Albright, T. D. (1984). Direction and orientation selectivity of neurons in visual area MT of the macaque. Journal of Neurophysiology, 52(6), 1106–1130. https://doi.org/10.1152/jn.1984.52.6.1106

Amunts, K., Malikovic, A., Mohlberg, H., Schormann, T., Zilles, K. (2000). Brodmann’s Areas 17 and 18 Brought into Stereotaxic Space—Where and How Variable? NeuroImage, 11(1), 66–84. https://doi.org/10.1006/nimg.1999.0516

Bae, G., & Luck, S. J. (2019). Decoding motion direction using the topography of sustained ERPs and alpha oscillations. NeuroImage, 184, 242–255. https://doi.org/10.1016/j.neuroimage.2018.09.029

Beck, J.M., Ma, W.J., Kiani, R., Hanks, T., Churchland, A.K., Roitman, J., Shadlen, M.N., Latham, P.E., Pouget, A. (2008). Probabilistic population codes for Bayesian decision making. Neuron 60(6). 1142–52. https://doi.org/10.1016/j.neuron.2008.09.021.

Bennur, S., & Gold, J. I. (2011). Distinct representations of a perceptual decision and the associated oculomotor plan in the monkey lateral intraparietal area. The Journal of Neuroscience: The Official Journal of the Society for Neuroscience, 31(3), 913–921. https://doi.org/10.1523/JNEUROSCI.4417-10.2011

Bode, S., Bogler, C., Haynes, J.-D. (2013). Similar neural mechanisms for perceptual guesses and free decisions. NeuroImage, 65, 456–465. https://doi.org/10.1016/j.neuroimage.2012.09.064

Braddick, O.J. (1973). The masking of apparent motion in random-dot patterns. Vision Research, 13(2), 355–369. https://doi.org/10.1016/0042-6989(73)90113-2

Braddick, O. J., O’Brien, J. M. D., Wattam-Bell, J., Atkinson, J., Hartley, T., & Turner, R. (2001). Brain Areas Sensitive to Coherent Visual Motion. Perception, 30(1), 61–72. https://doi.org/10.1068/p3048

Brainard, D. H., (1997). The Psychophysics Toolbox. Spatial Vision, 10(4), 433–436. https://doi.org/10.1163/156856897X00357

Brincat, S. L., Siegel, M., von Nicolai, C., Miller, E. K. (2018). Gradual progression from sensory to task-related processing in cerebral cortex. Proceedings of the National Academy of Sciences, 115(30), E7202–E7211. https://doi.org/10.1073/pnas.1717075115

Britten, K. H., Newsome, W. T., Shadlen, M. N., Celebrini, S., Movshon, J. A. (1996). A relationship between behavioral choice and the visual responses of neurons in macaque MT. Visual Neuroscience, 13(1), 87–100. https://doi.org/10.1017/S095252380000715X

Brodersen, K. H., Ong, C. S., Stephan, K. E., & Buhmann, J. M. (2010). The Balanced Accuracy and Its Posterior Distribution. 2010 20th International Conference on Pattern Recognition, 3121–3124. https://doi.org/10.1109/ICPR.2010.764

Brouwer, G. J., & Heeger, D. J. (2009). Decoding and reconstructing color from responses in human visual cortex. The Journal of Neuroscience, 29(44), 13992–14003. https://doi.org/10.1523/JNEUROSCI.3577-09.2009

Brouwer, G. J., & van Ee, R. (2007). Visual cortex allows prediction of perceptual states during ambiguous structure-from-motion. Journal of Neuroscience, 27(5), 1015–1023. https://doi.org/10.1523/JNEUROSCI.4593-06.2007

Caspers, S., Geyer, S., Schleicher, A., Mohlberg, H., Amunts, K., Zilles, K. (2006). The human inferior parietal cortex: Cytoarchitectonic parcellation and interindividual variability. NeuroImage, 33(2), 430–448. https://doi.org/10.1016/j.neuroimage.2006.06.054

Caywood, M.S., Roberts, D.M., Colombe, J.B., Greenwald, H.S. and Weiland, M.Z. (2017) Gaussian Process Regression for Predictive But Interpretable Machine Learning Models: An Example of Predicting Mental Workload across Tasks. Frontiers in Human Neuroscience, 10, 647. https://doi.org/10.3389/fnhum.2016.00647

Cichy, R.M., Heinzle, J., Haynes, J.D. (2012). Imagery and Perception Share Cortical Representations of Content and Location, Cerebral Cortex, 22(2), 372–380. https://doi.org/10.1093/cercor/bhr106

Chicharro, D., Panzeri, S., Haefner, R. M. (2021). Stimulus-dependent relationships between behavioral choice and sensory neural responses. eLife, 10, e54858. https://doi.org/10.7554/eLife.54858

Choi, H.-J., Zilles, K., Mohlberg, H., Schleicher, A., Fink, G. R., Armstrong, E., Amunts, K. (2006). Cytoarchitectonic identification and probabilistic mapping of two distinct areas within the anterior ventral bank of the human intraparietal sulcus. The Journal of Comparative Neurology, 495(1), 53–69. https://doi.org/10.1002/cne.20849

Churchland, A. K., Kiani, R., Shadlen, M. N. (2008). Decision-making with multiple alternatives. Nature Neuroscience, 11(6), 693–702. https://doi.org/10.1038/nn.2123

Cohen, B., Matsuo, V., Raphan, T.(1977). Quantitative analysis of the velocity characteristics of optokinetic nystagmus and optokinetic after-nystagmus. The Journal of Physiology, 270. https://doi.org/10.1113/jphysiol.1977.sp011955.

Christophel, T. B., Klink, P. C., Spitzer, B., Roelfsema, P. R., & Haynes, J. D. (2017). The Distributed Nature of Working Memory. Trends in cognitive sciences, 21(2), 111–124. https://doi.org/10.1016/j.tics.2016.12.007

Dimitrova, R., Pietsch, M., Christiaens, D., Ciarrusta, J., Wolfers, T., Batalle, D., Hughes, E., Hutter, J., Cordero-Grande, L., Price, A. N., Chew, A., Falconer, S., Vecchiato, K., Steinweg, J. K., Carney, O., Rutherford, M. A., Tournier, J. D., Counsell, S. J., Marquand, A. F., Rueckert, D., … O’Muircheartaigh, J. (2020). Heterogeneity in Brain Microstructural Development Following Preterm Birth. Cerebral cortex, 30(9), 4800–4810. https://doi.org/10.1093/cercor/bhaa069

Downing, C. J., & Movshon, J. A. (1989). Spatial and temporal summation in the detection of motion in stochastic random dot displays. Investigative Ophthalmology and Visual Science, 30, 72.

Dumoulin, S. O., & Wandell, B. A. (2008). Population receptive field estimates in human visual cortex. NeuroImage, 39(2), 647–660. 10.1016/j.neuroimage.2007.09.034

Eickhoff, S. B., Stephan, K. E., Mohlberg, H., Grefkes, C., Fink, G. R., Amunts, K., Zilles, K. (2005). A new SPM toolbox for combining probabilistic cytoarchitectonic maps and functional imaging data. NeuroImage, 25(4), 1325–1335. https://doi.org/10.1016/j.neuroimage.2004.12.034

Filimon, F., Philiastides, M. G., Nelson, J. D., Kloosterman, N. A., Heekeren, H. R. (2013). How Embodied Is Perceptual Decision Making? Evidence for Separate Processing of Perceptual and Motor Decisions. Journal of Neuroscience, 33(5), 2121–2136. https://doi.org/10.1523/JNEUROSCI.2334-12.2013

Forstmann, B. U., Ratcliff, R., Wagenmakers, E.J. (2016). Sequential Sampling Models in Cognitive Neuroscience: Advantages, Applications, and Extensions. Annual Review of Psychology, 67, 641–666. https://doi.org/10.1146/annurev-psych-122414-033645

Freedman, D. J., & Assad, J. A. (2011). A proposed common neural mechanism for categorization and perceptual decisions. Nature Neuroscience, 14(2), 143–146. https://doi.org/10.1038/nn.2740

Fusi, S., Miller, E. K., Rigotti, M. (2016). Why neurons mix: High dimensionality for higher cognition. Current Opinion in Neurobiology, 37, 66–74. https://doi.org/10.1016/j.conb.2016.01.010

Gardner, J. L., & Liu, T. (2019). Inverted Encoding Models Reconstruct an Arbitrary Model Response, Not the Stimulus. eNeuro, 6(2), ENEURO.0363-18.2019. https://doi.org/10.1523/ENEURO.0363-18.2019

Geisler, W. S. (1999). Motion streaks provide a spatial code for motion direction. Nature, 400(6739), 65–69. https://doi.org/10.1038/21886

Glasser, M. F., Sotiropoulos, S. N., Wilson, J. A., Coalson, T. S., Fischl, B., Andersson, J. L., Xu, J., Jbabdi, S., Webster, M., Polimeni, J. R., Van Essen, D. C., Jenkinson, M., WU-Minn HCP Consortium. (2013). The minimal preprocessing pipelines for the Human Connectome Project. NeuroImage, 80, 105–124. https://doi.org/10.1016/j.neuroimage.2013.04.127

Gold, J. I., & Shadlen, M. N. (2007). The Neural Basis of Decision Making. Annual Review of Neuroscience, 30(1), 535–574. https://doi.org/10.1146/annurev.neuro.29.051605.113038

Grind, W. A. van de, Doorn, A. J. van, Koenderink, J. J. (1983). Detection of coherent movement in peripherally viewed random-dot patterns. Journal of the Optical Society of America, 73(12), 1674–1683. https://doi.org/10.1364/JOSA.73.001674

Hanks, T. D., & Summerfield, C. (2017). Perceptual Decision Making in Rodents, Monkeys, and Humans. Neuron, 93(1), 15–31. https://doi.org/10.1016/j.neuron.2016.12.003

Haynes, J.D. (2015). A Primer on Pattern-Based Approaches to fMRI: Principles, Pitfalls, and Perspectives. Neuron, 87(2), 257–270. https://doi.org/10.1016/j.neuron.2015.05.025

Hebart, M. N., Donner, T. H., Haynes, J.D. (2012). Human visual and parietal cortex encode visual choices independent of motor plans. NeuroImage, 63(3), 1393–1403. https://doi.org/10.1016/j.neuroimage.2012.08.027

Hebart, M. N., Schriever, Y., Donner, T. H., Haynes, J.D. (2016). The Relationship between Perceptual Decision Variables and Confidence in the Human Brain. Cerebral Cortex, 26(1), 118–130. https://doi.org/10.1093/cercor/bhu181

Heekeren, H. R., Marrett, S., Bandettini, P. A., Ungerleider, L. G. (2004). A general mechanism for perceptual decision-making in the human brain. Nature, 431(7010), 859–862. https://doi.org/10.1038/nature02966

Heekeren, H. R., Marrett, S., Ruff, D. A., Bandettini, P. A., Ungerleider, L. G. (2006). Involvement of human left dorsolateral prefrontal cortex in perceptual decision making is independent of response modality. Proceedings of the National Academy of Sciences, 103(26), 10023–10028. https://doi.org/10.1073/pnas.0603949103

Heekeren, H. R., Marrett, S., Ungerleider, L.G. (2008). The neural systems that mediate human perceptual decision making. Nature Reviews Neuroscience, 9, 467–479. https://doi.org/10.1038/nrn2374

Ho, T. C., Brown, S., Serences, J. T. (2009). Domain General Mechanisms of Perceptual Decision Making in Human Cortex. The Journal of Neuroscience, 29(27), 8675–8687. https://doi.org/10.1523/JNEUROSCI.5984-08.2009

Huk, A. C., & Meister, M. L. (2012). Neural correlates and neural computations in posterior parietal cortex during perceptual decision-making. Frontiers in integrative neuroscience, 6, 86. https://doi.org/10.3389/fnint.2012.00086

Hummel, N., Cuturi, L. F., MacNeilage, P. R., Flanagin, V. L. (2016a). The effect of supine body position on human heading perception. Journal of Vision, 16(3), 19. https://doi.org/10.1167/16.3.19

Japkowicz, N., & Stephen, S. (2002). The class imbalance problem: A systematic study. Intelligent Data Analysis, 6(5), 429–449. http://dx.doi.org/10.3233/IDA-2002-6504

Kamitani, Y., & Tong, F. (2006). Decoding seen and attended motion directions from activity in the human visual cortex. Current biology, 16(11), 1096–1102. https://doi.org/10.1016/j.cub.2006.04.003

Kaplan, J. T., Man, K., Greening, S. G. (2015). Multivariate cross-classification: applying machine learning techniques to characterize abstraction in neural representations. Frontiers in Human Neuroscience. 9, 151. https://doi.org/10.3389/fnhum.2015.00151

Kay, K. N., Naselaris, T., Prenger, R. J., Gallant, J. L. (2008). Identifying natural images from human brain activity. Nature, 452(7185), 352–355. https://doi.org/10.1038/nature06713

Kleiner, M., Brainard, D., Pelli, D. (2007). What’s new in Psychtoolbox-3?. Perception 36 ECVP Abstract Supplement.

Kriegeskorte, N., Cusack, R., & Bandettini, P. (2010). How does an fMRI voxel sample the neuronal activity pattern: compact-kernel or complex spatiotemporal filter?. NeuroImage, 49(3), 1965–1976. https://doi.org/10.1016/j.neuroimage.2009.09.059

Kriegeskorte, N., & Diedrichsen, J. (2019). Peeling the Onion of Brain Representations. Annual Review of Neuroscience, 42(1), 407–432. https://doi.org/10.1146/annurev-neuro-080317-061906

Kriegeskorte, N., Goebel, R., Bandettini, P. (2006). Information-based functional brain mapping. Proceedings of the National Academy of Sciences, 103(10), 3863–3868. https://doi.org/10.1073/pnas.0600244103

Krishna, A., Tanabe, S., Kohn, A. (2021). Decision signals in the local field potentials of early and mid-level macaque visual cortex. Cerebral Cortex, 31(1), 169–183. https://doi.org/10.1093/cercor/bhaa218

Kujovic, M., Zilles, K., Malikovic, A., Schleicher, A., Mohlberg, H., Rottschy, C., Eickhoff, S. B., Amunts, K. (2013). Cytoarchitectonic mapping of the human dorsal extrastriate cortex. Brain Structure and Function, 218(1), 157–172. https://doi.org/10.1007/s00429-012-0390-9

Levine, S. M., & Schwarzbach, J. (2017). Decoding of auditory and tactile perceptual decisions in parietal cortex. NeuroImage, 162, 297–305. https://doi.org/10.1016/j.neuroimage.2017.08.060

Levinson, E., & Sekuler, R. (1976). Adaptation alters perceived direction of motion. Vision Research, 16(7), 779-IN7. https://doi.org/10.1016/0042-6989(76)90189-9

Liu, T., & Pleskac, T. J. (2011). Neural correlates of evidence accumulation in a perceptual decision task. Journal of Neurophysiology, 106(5), 2383–2398. https://doi.org/10.1152/jn.00413.2011

Maris, E., Oostenveld, R. (2007). Nonparametric statistical testing of EEG and MEG-data. Journal of Neuroscience Methods, 164(1), 177–190. https://doi.org/10.1016/j.jneumeth.2007.03.024

Merriam, E.P., Gardner, J.L., Movshon, J.A., Heeger, D.J. (2013). Modulation of visual responses by gaze direction in human visual cortex. The Journal of Neuroscience, 33(24), 9879–9889. https://doi.org/10.1523/JNEUROSCI.0500-12.2013

Movshon, J. A., & Newsome, W. T. (1996). Visual response properties of striate cortical neurons projecting to area MT in macaque monkeys. Journal of Neuroscience, 16(23), 7733–7741. https://doi.org/10.1523/JNEUROSCI.16-23-07733.1996

Mulder M.J., van Maanen L., Forstmann B.U. (2014). Perceptual decision neurosciences—a model-based review. Neuroscience. 2014; 277:872–84. https://doi.org/10.1016/j.neuroscience.2014.07.031

Naselaris T., Kay K.N., Nishimoto S., Gallant J.L. (2011). Encoding and decoding in fMRI. NeuroImage, 56, 400–410. https://doi.org/10.1016/j.neuroimage.2010.07.073

Nevado, A., Young, M. P., Panzeri, S. (2004). Functional imaging and neural information coding. NeuroImage, 21(3), 1083–1095. https://doi.org/10.1016/j.neuroimage.2003.10.043

Newsome, W. T., & Paré, E. B. (1988). A selective impairment of motion perception following lesions of the middle temporal visual area (MT). The Journal of Neuroscience, 8(6), 2201– 2211. https://doi.org/10.1523/jneurosci.08-06-02201.1988

Nichols, M. J., & Newsome, W. T. (2002). Middle temporal visual area microstimulation influences veridical judgments of motion direction. The Journal of Neuroscience, 22(21), 9530–9540. https://doi.org/10.1523/JNEUROSCI.22-21-09530.2002

Park, I., Meister, M., Huk, A., Pillow, J.W. (2014). Encoding and decoding in parietal cortex during sensorimotor decision-making. Nature Neuroscience 17, 1395–1403. https://doi.org/10.1038/nn.3800

Pilly, P. K., & Seitz, A. R. (2009). What a difference a parameter makes: A psychophysical comparison of random dot motion algorithms. Vision Research, 49(13), 1599–1612. https://doi.org/10.1016/j.visres.2009.03.019

Pratte, M.S., and Tong, F. (2014). Spatial specificity of working memory representations in the early visual cortex. Journal of Vision. 14, 22. http://dx.doi.org/10.1167/14.3.22

Prinzmetal, W., Amiri, H., Allen, K., Edwards, T. (1998). Phenomenology of attention: I. Color, location, orientation, and spatial frequency. Journal of Experimental Psychology: Human Perception and Performance, 24(1), 261. https://doi.org/10.1037/0096-1523.24.1.261

Ramírez, F. M., Cichy, R. M., Allefeld, C., & Haynes, J. D. (2014). The neural code for face orientation in the human fusiform face area. The Journal of Neuroscience, 34(36), 12155– 12167. https://doi.org/10.1523/JNEUROSCI.3156-13.2014

Rasmussen, C. E., & Nickisch, H. (2010). Gaussian processes for machine learning (GPML) toolbox. The Journal of Machine Learning Research, 11, 3011–3015.

Rasmussen, C. E., & Williams, C. K. I. (2005). Gaussian processes for machine learning. MIT Press. https://doi.org/10.7551/mitpress/3206.001.0001

Ratcliff R. (2018). Decision making on spatially continuous scales. Psychological review, 125(6), 888–935. https://doi.org/10.1037/rev0000117

Rees, G., Friston, K., Koch, C. (2000). A direct quantitative realtionship between the functional properties of human and macaque V5. Nature Neuroscience, 3, 716–723. https://doi.org/10.1038/76673

Ress, D., Heeger, D.J. (2003). Neuronal correlates of perception in early visual cortex. Nature Neuroscience, 6(4), 414–20. https://doi.org/10.1038/nn1024

Riggall, A. C., & Postle, B. R. (2012). The relationship between working memory storage and elevated activity as measured with functional magnetic resonance imaging. The Journal of Neuroscience, 32(38), 12990–12998. https://doi.org/10.1523/JNEUROSCI.1892-12.2012

Rigotti, M., Barak, O., Warden, M. R., Wang, X.-J., Daw, N. D., Miller, E. K., Fusi, S. (2013). The importance of mixed selectivity in complex cognitive tasks. Nature, 497(7451), 585–590. https://doi.org/10.1038/nature12160

Rissman, J., Gazzaley, A., D’Esposito, M. (2004). Measuring functional connectivity during distinct stages of a cognitive task. Neuroimage, 23(2), 752–763. https://doi.org/10.1016/j.neuroimage.2004.06.035

Rottschy, C., Eickhoff, S. B., Schleicher, A., Mohlberg, H., Kujovic, M., Zilles, K., Amunts, K. (2007). Ventral visual cortex in humans: Cytoarchitectonic mapping of two extrastriate areas. Human Brain Mapping, 28(10), 1045–1059. https://doi.org/10.1002/hbm.20348

Scheperjans, F., Eickhoff, S. B., Hömke, L., Mohlberg, H., Hermann, K., Amunts, K., Zilles, K. (2008). Probabilistic Maps, Morphometry, and Variability of Cytoarchitectonic Areas in the Human Superior Parietal Cortex. Cerebral Cortex, 18(9), 2141–2157. https://doi.org/10.1093/cercor/bhm241

Serences, J. T., & Boynton, G. M. (2007). The Representation of Behavioral Choice for Motion in Human Visual Cortex. Journal of Neuroscience, 27(47), 12893–12899. https://doi.org/10.1523/JNEUROSCI.4021-07.2007

Serences, J. T., Ester, E. F., Vogel, E. K., & Awh, E. (2009). Stimulus-specific delay activity in human primary visual cortex. Psychological science, 20(2), 207–214. https://doi.org/10.1111/j.1467-9280.2009.02276.x

Shadlen M.N., Britten K.H., Newsome W.T., Movshon J.A. (1996). A computational analysis of the relationship between neuronal and behavioral responses to visual motion. Journal of Neuroscience, 16, 1486–510. https://doi.org/10.1523/JNEUROSCI.16-04-01486.1996

Shadlen, M. N., Kiani, R., Hanks, T. D., Churchland, A. K. (2008). Neurobiology of decision making: An intentional framework. In C. Engel & W. Singer (Eds.), Better than conscious? Decision making, the human mind, and implications for institutions (pp. 71–101). MIT Press. https://doi.org/10.7551/mitpress/9780262195805.003.0004

Siegel, M., Engel, A. K., Donner, T. H. (2011). Cortical network dynamics of perceptual decision-making in the human brain. Frontiers in Human Neuroscience, 5, 21. https://doi.org/10.3389/fnhum.2011.00021

Smith, P. L. (2016). Diffusion theory of decision making in continuous report. Psychological Review, 123(4), 425–451. https://doi.org/10.1037/rev0000023

Sousa, T., Duarte, J.V., Costa, G.N., Kemper, V.G., Martins, R., Goebel, R., Castelo-Branco, M. (2021). The dual nature of the BOLD signal: Responses in visual area hMT+ reflect both input properties and perceptual decision. Human Brain Mapping. 42, 1920– 1929. https://doi.org/10.1002/hbm.25339

Sprague, T. C., Adam, K., Foster, J. J., Rahmati, M., Sutterer, D. W., & Vo, V. A. (2018). Inverted Encoding Models Assay Population-Level Stimulus Representations, Not Single-Unit Neural Tuning. eNeuro, 5(3), ENEURO.0098-18.2018. https://doi.org/10.1523/ENEURO.0098-18.2018

Sprague, T. C., Boynton, G. M., & Serences, J. T. (2019). The Importance of Considering Model Choices When Interpreting Results in Computational Neuroimaging. eNeuro, 6(6), ENEURO.0196-19.2019. https://doi.org/10.1523/ENEURO.0196-19.2019

Thaler, L., Schütz, A. C., Goodale, M. A., Gegenfurtner, K. R. (2013). What is the best fixation target? The effect of target shape on stability of fixational eye movements. Vision Research, 76, 31–42. https://doi.org/10.1016/j.visres.2012.10.012

Thielen, J., Bosch, S.E., van Leeuwen, T.M., van Gerven, M.A.J., van Lier, R. (2019). Evidence for confounding eye movements under attempted fixation and active viewing in cognitive neuroscience. Scientific Reports, 9, 17456. https://doi.org/10.1038/s41598-019-54018-z

Thielen, J., van Lier, R., & van Gerven, M. (2018). No evidence for confounding orientation-dependent fixational eye movements under baseline conditions. Scientific Reports, 8(1), 1–10. https://doi.org/10.1038/s41598-018-30221-2

Thirion, B., Duchesnay, E., Hubbard, E., Dubois, J., Poline, J.-B., Lebihan, D., Dehaene, S. (2006). Inverse retinotopy: inferring the visual content of images from brain activation patterns. NeuroImage, 33(4), 1104–1116. https://doi.org/10.1016/j.neuroimage.2006.06.062

Töpfer, F. M., Barbieri, R., Sexton, C. M., Wang, X., Soch, J., Bogler, C., & Haynes, J. D. (2022). Psychophysics and computational modeling of feature-continuous motion perception. Journal of Vision, 22(11), 16. https://doi.org/10.1167/jov.22.11.16

Tosoni, A., Corbetta, M., Calluso, C., Committeri, G., Pezzulo, G., Romani, G. L., Galati, G. (2014). Decision and action planning signals in human posterior parietal cortex during delayed perceptual choices. European Journal of Neuroscience, 39(8), 1370–1383. https://doi.org/10.1111/ejn.12511

Tosoni, A., Galati, G., Romani, G. L., Corbetta, M. (2008). Sensory-motor mechanisms in human parietal cortex underlie arbitrary visual decisions. Nature Neuroscience, 11(12), 1446– 1453. https://doi.org/10.1038/nn.2221

Uchida, N., Kepecs, A., Mainen, Z. (2006). Seeing at a glance, smelling in a whiff: rapid forms of perceptual decision making. Nature Reviews Neuroscience, 7, 485–491. https://doi.org/10.1038/nrn1933

Urai, A., Braun, A., Donner, T. (2017). Pupil-linked arousal is driven by decision uncertainty and alters serial choice bias. Nature Communications 8, 14637. https://doi.org/10.1038/ncomms14637

van Bergen, R. S., Ma, W. J., Pratte, M. S., Jehee, J. F. (2015). Sensory uncertainty decoded from visual cortex predicts behavior. Nature Neuroscience, 18(12), 1728–1730. https://doi.org/10.1038/nn.4150

Wang, H. X., Merriam, E. P., Freeman, J., & Heeger, D. J. (2014). Motion direction biases and decoding in human visual cortex. The Journal of Neuroscience, 34(37), 12601–12615. https://doi.org/10.1523/JNEUROSCI.1034-14.2014

Watson, A. B., & Pelli, D. G. (1983). QUEST: A Bayesian adaptive psychometric method. Perception & psychophysics, 33(2), 113–120. https://doi.org/10.3758/BF03202828

Wilbertz, G., Ketkar, M., Guggenmos, M., Sterzer, P. (2018). Combined fMRI-and eye movement-based decoding of bistable plaid motion perception. Neuroimage, 171, 190–198. https://doi.org/10.1016/j.neuroimage.2017.12.094

Wilming, N., Murphy, P.R., Meyniel, F., Donner, T.H. (2020). Large-scale dynamics of perceptual decision information across human cortex. Nature Communications 11, 5109. https://doi.org/10.1038/s41467-020-18826-6

Zhang, W., & Luck, S. J. (2008). Discrete fixed-resolution representations in visual working memory. Nature, 453(7192), 233–235. https://doi.org/10.1038/nature06860

